# Precision editing of an aggression-encoding network relay suppresses violent action in mice

**DOI:** 10.1101/2022.12.07.519272

**Authors:** Yael S. Grossman, Austin Talbot, Neil M. Gallagher, Kathryn K. Walder-Christensen, Gwenaëlle E. Thomas, Alexandra Fink Skular, Scott J. Russo, David E. Carlson, Kafui Dzirasa

**Affiliations:** Howard Hughes Medical Institute, Chevy Chase, Maryland 20815, USA; Dept. of Psychiatry and Behavioral Sciences, Duke University Medical Center, Durham, North Carolina 27710, USA; Department of Statistical Science, Duke University, Durham North Carolina 27708, USA; Dept. of Neurobiology, Duke University, Durham North Carolina 27708, USA; Fishberg Department of Neuroscience and Friedman Brain Institute, Icahn School of Medicine at Mount Sinai, New York, New York, 10029, USA; Dept. of Biostatistics and Bioinformatics, Duke University Medical Center, Durham, North Carolina 27710, USA; Dept. of Civil and Environmental Engineering; Dept. of Biomedical Engineering, Duke University, Durham North Carolina 27708, USA; Dept. of Neurosurgery, Duke University Medical Center, Durham, North Carolina 27710, USA

**Keywords:** aggression, social behavior, electome network, closed-loop stimulation, machine learning, medial prefrontal cortex

## Abstract

Aggression is a psychological state characterized by intent to harm oneself or others, and often manifests as hostile, harmful, or violent action. Here we showed that an aggressive internal state in mice is encoded by a widespread brain network. This network is organized by prominent and synchronized electrical oscillations that relay through the medial prefrontal cortex, and network activity is coupled to widespread cellular firing. Strikingly, network activity during social isolation encoded the innate aggressiveness of mice, and causal cellular manipulations known to impact aggression modulated the network’s activity. We established that this network mediates aggression using closed-loop stimulation of medial prefrontal cortex and causal mediation analysis. Finally, we applied ’Long-term Integration of Circuits using connexins’ (LinCx) to selectively edit a key circuit within the network from the medial prefrontal cortex to the nucleus accumbens. Editing this circuit durably suppressed violent action while leaving non-aggressive social behavior intact.

## INTRODUCTION

Social aggression can aid an organism in securing access to resources^1, 2^, or it can impair group function and survival in behavioral pathology^3, 4, 5^. Since many brain regions contribute to multiple social behaviors^6, 7, 8^, expanded knowledge of how the brain distinguishes between social states would enable the development of interventions that suppress violent action, while leaving other social behaviors intact. Indeed, social behavior reflects the integration of sensory information with internal affective states. Many subcortical brain regions contribute to aggressive behavior in mammals including lateral septum (LSN) ^9, 10^, nucleus accumbens (NAc)^3, 11^, lateral habenula (LHb) ^12, 13^, the ventrolateral portion of ventromedial hypothalamus (VMHvl) ^6, 14, 15, 16, 17, 18^, and medial amygdala (MeA) ^4, 19^. Medial prefrontal cortex stimulation has been shown to mitigate aggressive behavior in both humans ^20, 21^ and rodents ^22^, implicating cortical regions in regulating aggression. Finally, sensory regions, such as those underlying olfaction^23, 24^, also contribute to aggression.

To appropriately regulate aggressive behavior, the brain must integrate information across these and other cortical and subcortical regions, many of which also regulate non-aggressive social behaviors ^6, 25^. Therefore, the brain must integrate information from overlapping regions to distinguish aggression from other social behavioral states. Though efforts have revealed several cellular-level processes within these regions that contribute to this mechanism ^14, 17, 26^, the complementary network-level process that integrates information across regions to distinguish aggressive states from pro-social states has not been characterized. Addressing this knowledge gap is of major importance as 1) mammals regularly select from a repertoire of social behaviors based on external sensory cues to ensure their survival ^7, 27^, and 2) a range of psychiatric disorders are broadly marked by a failure to appropriately match behavior with evolving social contexts ^28^. Because independent changes arising from many distinct regions may potentially converge to alter aggression regulation, the optimal circuit to target for widescale therapeutic intervention remains unclear.

## RESULTS

### Direct stimulation of medial prefrontal cortex broadly suppresses social behaviors

We initially focused on modulating the medial prefrontal cortex (mPFC; i.e., prelimbic and infralimbic cortex in mice) as a means to selectively suppress aggression, since this brain region had been implicated in social behaviors ^29, 30, 31^. Moreover, clinical studies have shown that direct transcranial stimulation of mPFC decreased aggressive feelings in violent offenders ^32^ and individuals with methamphetamine use disorder ^33^. Thus, we performed optogenetic stimulation of mPFC during social encounters using a protocol that was modeled after prior work (Fig. 1a)^22^.

**Figure 1.**
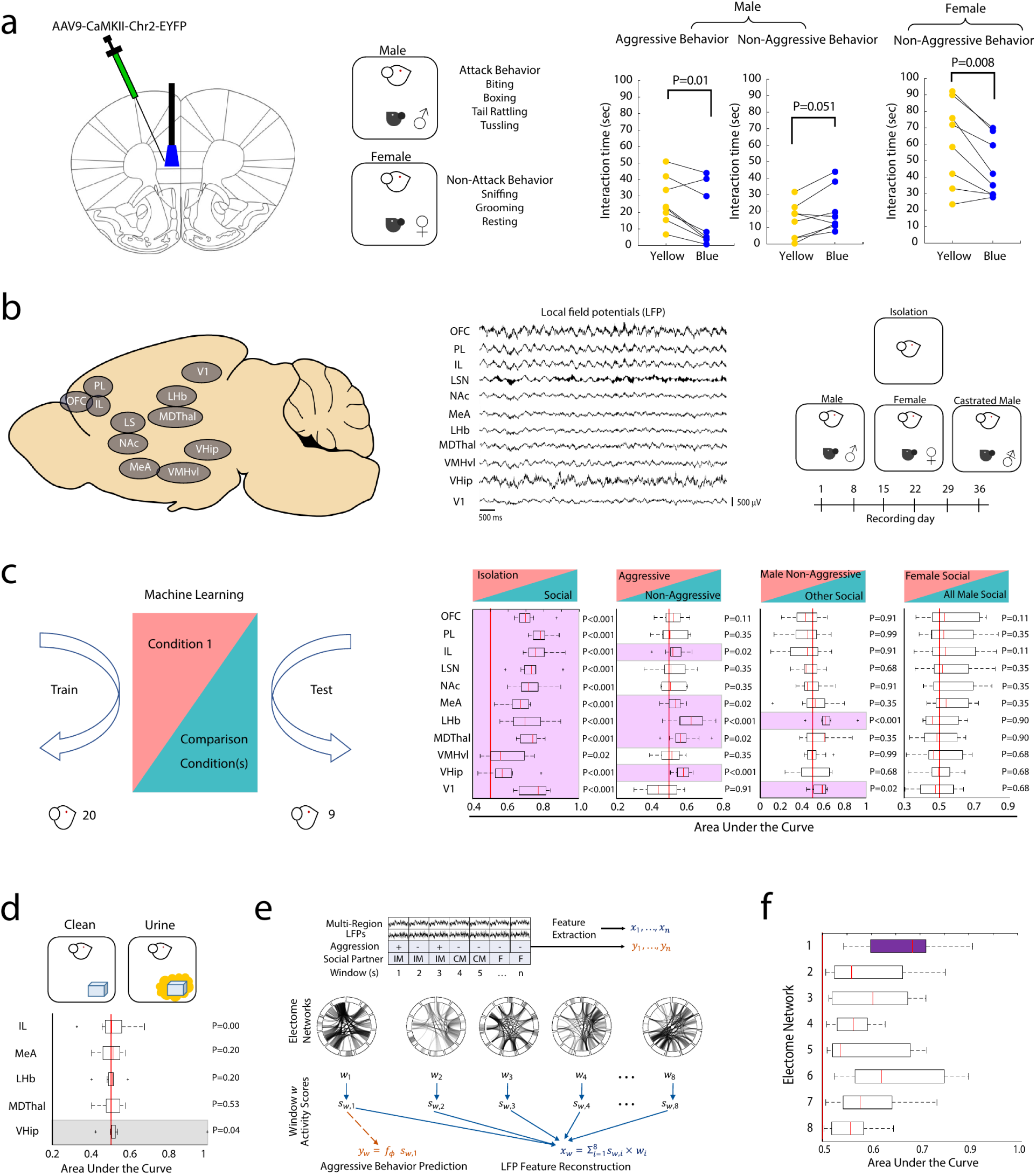
A widespread network encodes aggressive behavior. **a)** Direct stimulation of medial prefrontal cortex suppresses social behavior. Schematic of optogenetic stimulation (left) and social encounters utilized for testing (middle). Right: Medial prefrontal cortex stimulation suppressed aggressive behavior, increased non-aggressive social behavior towards male mice, and suppressed non-aggressive social behavior towards females (stimulation using blue - 473nm vs. yellow - 593.5nm light, 5mW, 20Hz, 3ms pulse width, P<0.05 for each comparison using two-tailed paired t-tests, n=8). **b)** Schematic for electrical recordings, showing targeted brain regions (left) and representative local field potentials (middle) recorded on the days indicated on the bottom timeline during isolation in the home cage, as well as during repeated exposure to social contexts that produce aggressive (e.g., exposure to an intact male) and non-aggressive (e.g., exposure to female or castrated male mouse) social behavior (right). **c)** Framework to test individual brain regions’ encoding of social states (left). All implanted regions distinguished social engagement from isolation in test mice (N=9, right, 1^st^ graph); however, only five selectively encoded the aggressive behavior vs. non-aggressive behavior (right, 2^nd^ graph). Pink shading indicates P<0.05 using one-tailed Wilcoxon rank-sum, significance determined by FDR correction for 44 comparisons. **d)** The most aggressive mice from the training group (N=8) were allowed to freely explore a clean inanimate object or an object covered in urine from another intact CD1 male mouse over seven sessions. Neural signatures for aggression discovered from the five individual brain regions in panel (c) failed to encode aggressive behavior induced by male urine (gray shading indicates P<0.05 using one-tailed Wilcoxon rank-sum prior to, but not following, FDR correction). **e)** Schematic of electome-based decoding and reconstruction of aggressive state from multi-region LFP activity. (Top) Multi-region LFP recordings are segmented into time windows aligned with labels of non-aggressive/aggressive behavior and the sex of the intruder. LFPs are transformed into features describing distributed neural activity. (Middle) These features and labels are then used to train a machine learning model that learns electome networks with corresponding network-level activity patterns across windows. (Bottom) The network activity from a single network is then used to predict aggressive behavior (left), while the learned network structure enables reconstruction of LFP features across brain regions (right), providing an interpretable representation of the distributed neural dynamics associated with aggression. **f)** Encoding across eight learned networks (1 supervised network and 7 unsupervised networks) in hold-out test mice (N=9). The supervised network (purple bar) showed the strongest encoding (AUC=0.67±0.03, one-tailed Wilcoxon rank sum, P=2×10^-5^). Data shown as |AUC-0.05| across the group for each network. Error bars represent ± SEM of the AUC across mice for each network. The red line corresponds with an AUC of 0.5, which represents the model’s inability to distinguish between aggressive and pro-social behavior. For the box and whisker plot: central mark is the median, box edges are the 25th and 75th percentiles, whiskers extend to extreme data points that are not outliers, and outliers are plotted individually as "+." Source data are provided as a Source Data file.

To test the behavioral effects of mPFC stimulation, we examined CD1 mice in two established social paradigms. When a male C57BL6/J (C57) mouse is introduced into their home cage, CD1 mice show periods of violent/aggressive behavior [biting, boxing (kicking/clawing), or tussling] ^34^. In contrast, when exposed to a female intruder, CD1 mice exhibit periods of non-aggressive social interactions [sniffing, grooming, or resting (placing nose or forepaws against the subject mouse, but not moving)] ^17^. Importantly, the CD1 mice do not show violent behavior during this latter social context.

Consistent with the prior reports, mPFC stimulation suppressed aggressive behavior towards C57 male mice intruders and tended to increase non-aggressive social behavior [N=8; t_7_=3.48; P=0.01, and t_7_=-2.35; P=0.051 using a two-tailed paired t-test, for aggressive and non-aggressive behavior, respectively, significance determined by a Benjamini-Hochberg false discovery rate (FDR) correction, Fig. 1a]. However, this suppression of social behavior towards intruders was not specific to aggressive behavior, as we found that mPFC stimulation decreased non-aggressive social behavior toward female intruders as well (t_7_=3.647, P=0.008, using two-tailed paired t-test, significance determined by FDR correction). Thus, mPFC stimulation suppressed multiple types of social behavior, not solely aggressive behavior.

### Individual brain regions fail to independently encode aggressive behavior across contexts

After failing to selectively suppress CD1 aggressive social behavior by targeting the mPFC, we set out to find brain regions that may exhibit such selectivity. We implanted CD1 mice across multiple cortical and subcortical brain regions known to contribute to social behavior, including infralimbic cortex^22^, orbitofrontal cortex^35^, prelimbic cortex^22^, lateral septum ^9, 10^, nucleus accumbens ^3, 11^, lateral habenula ^12, 13^, mediodorsal thalamus^36^, ventromedial hypothalamus ^6, 14, 15, 16, 17, 18^, medial amygdala ^4, 19^, ventral hippocampus ^37^, and primary visual cortex.

Following surgical recovery, we recorded neural activity while the CD1 mice freely interacted with an intact male C57 mouse and a female C57 mouse for 300 seconds each. We repeated these encounters over six sessions, yielding a total of 1800 seconds of neural data and behavior for each exposure (Fig. 1b, see also Supplementary Fig. S1). We recorded a subset of these CD1 mice (N=9 of the 20 implanted mice) as they interacted with a castrated male mouse intruder. Since CD1 mice do not generally exhibit aggressive behavior towards castrated males ^24^, this encounter provided neurophysiology data during additional non-aggressive social behaviors that were not linked to female sensory cues. We also acquired neural activity while the CD1 mice were isolated in their home cage.

We first verified that each of the implanted brain regions encoded social behavior using discriminative cross spectral factor analysis - nonnegative matrix factorization (dCSFA-NMF, see Fig. 1e) ^38^. dCSFA-NMF is a statistical model that integrates brain local field potential (LFP) activity features across time and discovers networks that distinguish between types of external behavior. LFPs reflect the activity of populations of neurons, and these signals can be consistently sampled across mice. The electrical functional connectome networks (electome networks) learned in dCSFA-NMF integrate LFP power (oscillatory amplitude across 1-56 Hz; a correlate of cellular and synaptic activity within brain regions), coherence (how two regions’ LFP synchronize across time; a correlate of brain circuit function), and directionality (assessed via Granger causality testing; a correlate of directional transfer of information across a brain circuit). Furthermore, dCSFA-NMF produces electome network activity scores, which indicate the strength of each network at a temporal resolution sufficient to capture brain states underlying relevant external behavior (in this instance, a resolution of one second). Any given brain region can belong to multiple electome networks, and each electome network may incorporate any number of brain regions. dCSFA-NMF thus integrates spatially distinct brain regions and circuits into interpretable networks that encode measured behavior^39^.

To explore whether there was a generalized activity pattern within individual regions that encoded social behavior, we fit dCSFA-NMF models based solely on 1-56Hz LFP oscillatory power in a single region. Each single-region model was trained using data pooled from 20 CD1 mice to separate periods when mice were socially isolated from periods when they were engaged in social behavior (e.g., aggressive behavior towards the intact males and non-aggressive social behavior towards intact and castrated male mice and female mice). We trained our models with one supervised network to discover the patterns of LFP activity that encoded social behavior, meaning that this one network was encouraged to relate to the behaviors of interest and the LFP features. We also trained three unsupervised networks that were encouraged to only explain the variance in LFP features (see methods for justification of hyperparameter selection). We tested the accuracy of these models in new hold-out CD1 mice exposed to intruder male and female mice (N=9 mice).

We observed high accuracy in distinguishing periods of isolation from social behavior, using the supervised network, for each implanted brain area (p<0.05 using one-tailed Wilcoxon rank-sum, significance determined by FDR correction for 44 comparisons, Fig. 1c, right, 1^st^ graph). Next, we designed a new series of models to separate one of the three social behavioral states from the other two (e.g., aggressive behavior toward intact males vs. non-aggressive interactions with female and non-aggressive interactions with intact/castrated male mice). These models were built using data pooled from the same 20 CD1 mice and were again based on LFP power. We then tested the accuracy of the supervised networks for these single-region models in decoding within the same nine hold-out mice. Thus, we trained and tested the model’s generalizability to decode each class of social behavior from the other two for each of the 11 implanted brain areas (i.e., 33 additional models). Using this approach, we found that five of the brain region-based statistical models decoded aggressive behavior versus non-aggressive social behavior: infralimbic cortex, medial amygdala, lateral habenula, medial dorsal thalamus and ventral hippocampus. V1 and lateral habenula successfully decoded the non-aggressive male interaction from the other social conditions as well (P<0.05 using one-tailed Wilcoxon rank-sum test, significance determined by FDR correction for 44 comparisons, see Fig. 1c, right, 2^nd^ graph). None of the other implanted brain regions showed selectivity in its encoding for one type of social behavior. Thus, five regions independently encoded violent action vs. non-aggressive social behavior in a manner that generalized across mice in this particular social context.

However, our overall goal was to identify a neural signature that could be used to suppress aggression across multiple contexts, while leaving other social behaviors intact. Thus, we tested whether these five putative single-region neural signatures would generalize to another context associated with aggression. Specifically, urine from other male mice has been found to elicit aggressive and dominance behavior in CD1 males ^23, 24, 40,41^. As such, the most aggressive mice from the training group (N=8 of the 20 training mice, see methods) were allowed to freely explore a clean inanimate object or an object covered in urine from another intact CD1 male mouse (seven sessions). We then tested whether the significant single-region models distinguished periods when mice explored clean objects from those when mice explored objects covered in urine. Though only the ventral hippocampus model tended to decode behavior in this new context, none of the brain regions showed statistically significant encoding at the mesoscale-level (LFP) following multiplicity correction (P>0.05 using one-tailed Wilcoxon rank-sum test, significance determined by FDR correction for 5 comparisons, Fig. 1d).

### Aggressive behavior is encoded at the network-level across mice and contexts

After failing to robustly decode aggressive behavior across contexts solely using LFPs from one of the 11 brain regions, we established that a brain network integrated information across all the implanted brain regions to encode aggressive behavior across mice. This network-level encoding mechanism generalized to multiple new contexts associated with aggression.

For this analysis, we trained a new dCSFA-NMF model using data from all the implanted brain regions. This model utilized LFP power for each region and the coherence and directionality between regions. The model utilized one supervised network that was trained to encode periods of aggressive behavior (positive class) vs. non-aggressive social behavior towards castrated males and the female social context (negative class), in addition to capturing variance in the neural activity. We included non-aggressive social behavior towards intact males in the negative class to discourage the network from simply learning sensory cues specific to the intact male. Based on our hyperparameter selection approach using the Bayesian Information Criteria (see methods and Supplementary Fig. S2), seven additional unsupervised networks were trained solely to account for the variance in neural activity. We then validated our supervised network using the same set of nine hold-out CD1 mice as our single-region analysis. Again, none of these mice were used to train the electome networks. We found that the supervised network successfully discriminated between aggressive behavior and non-aggressive social behavior in the test mice (electome network #1; N=9 hold-out test mice, AUC=0.67±0.03, one-tailed Wilcoxon rank sum against chance, P=2×10^-5^ , Fig. 1f and Supplementary Fig. S3-5). Activity in this supervised network was lower when animals exhibited aggressive behavior, while high activity in one of the unsupervised networks mapped directly to aggressive behavior [Network #6; AUC=0.65±0.03, P<5×10^-5^ using one-tailed Wilcoxon rank sum against chance (AUC=0.5, signifying the model’s inability to distinguish between aggressive behavior and pro-social behavior); 9 hold-out test mice, see Supplementary Fig. S3].

In contrast to the single-region models developed for each of the brain regions, electome network #1 activity also encoded the exposure to male urine (N=8, Network Activity = 9.1±1.0 and 8.3±1.1 for clean and urine covered objects, respectively; P=0.012 using one-tailed Wilcoxon signed-rank test to evaluate whether exposure to urine decreased network activity; AUC=0.57±0.02, data not shown). Electome network #6 activity failed to generalize to this urine context (AUC=0.48±0.01, data not shown). Thus, we focused our subsequent analyses on electome network #1.

### An aggressive psychological state encoded at the network-level

Aggressive behavior is indicative of the aggressive psychological state. Yet, we also reasoned that it was possible for a mouse to be in an aggressive psychological state without being actively engaged in aggressive behavior. Since such a context was likely to be present immediately prior to or following the expression of aggressive behavior, we tested whether network activity pooled from the 3 seconds preceding and 3 seconds following social behaviors encoded the distinct social conditions (Fig. 2a; N=17 mice from the training set). These data windows were not used to train the network model since they did not contain the behaviors of interest. Activity of the supervised network (electome network #1) was lower in the intervals surrounding aggressive behavior compared to isolation and both non-aggressive social interactions with males and females (F_3,67_=28.6, P=2.78×10^-6^, Friedman test; post-hoc analysis performed using two-tailed Wilcoxon signed-rank test, P<0.005 after FDR correction, Fig. 2b, left). Strikingly, electome network #1 activity during periods of isolation also negatively correlated with the time mice spent exhibiting aggressive behavior towards other males (R=-0.58; P=0.016 using Spearman’s rank correlation, Fig. 2b, right), encoding an aggressive psychological state on a mouse-by-mouse basis. For comparison, we also performed similar analysis for VHip, since this region had shown the largest AUC for the single-region models in the urine context (Fig. 1d). VHip failed to encode the aggressive psychological state during the intervals surrounding aggressive behavior, and VHip activity in the home cage also failed to encode individual behavior (see Supplementary Fig. S4). Thus, electome network #1 uniquely encoded aggressive behavior across multiple contexts and signaled a generalized aggressive internal state that extended beyond the moments of overt aggressive behavior. Since decreased electome network #1 activity corresponded with aggression, these results suggested the network [hereafter referred to as *EN-Aggression Inhibition (EN-AggINH*)] putatively inhibits aggression when it is active.

**Figure 2.**
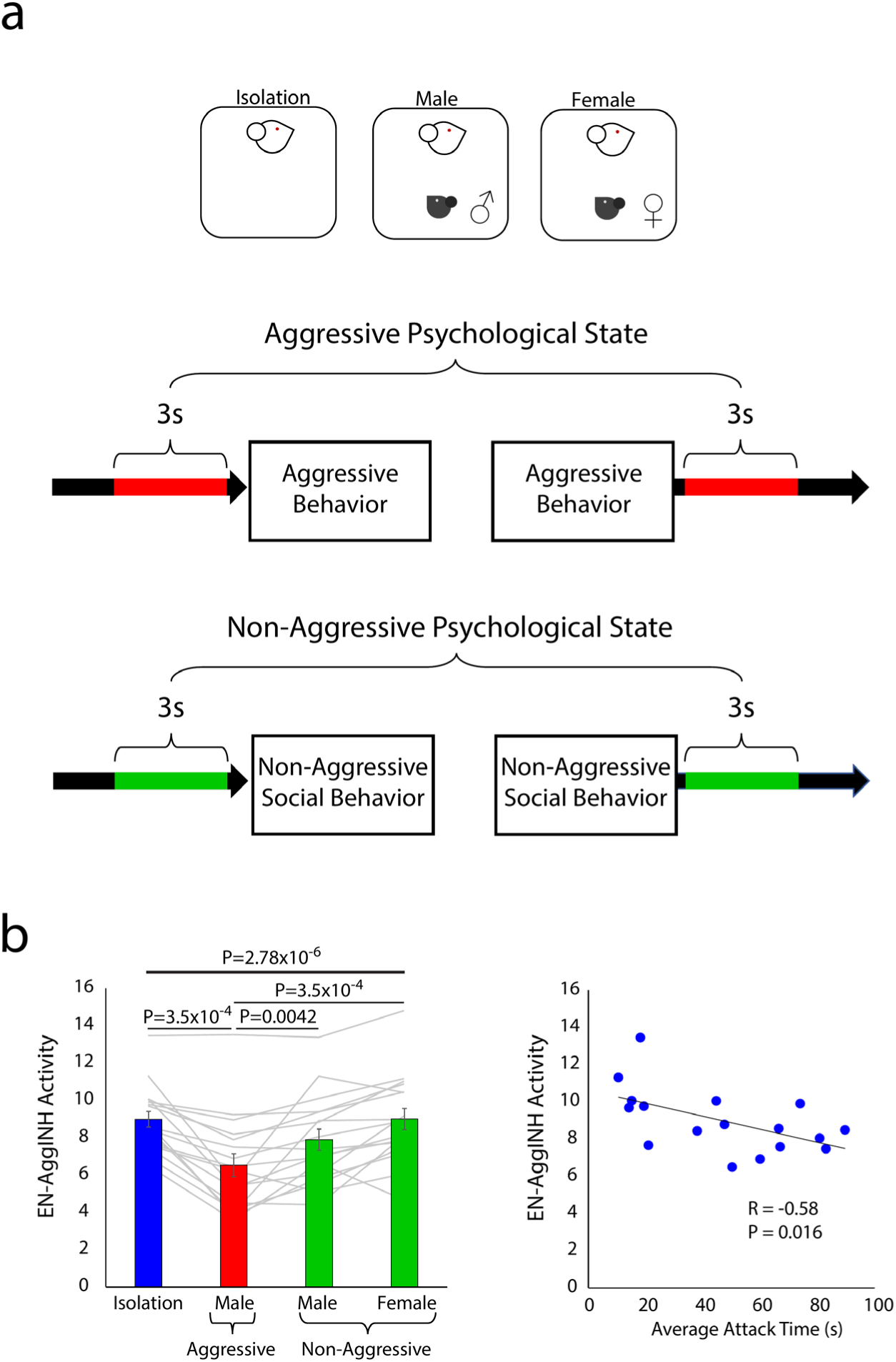
EN-Aggression Inhibition (EN-AggINH) encodes an aggressive internal state. **a)** Neural activity was sampled while 17 mice were socially isolated (blue) and during intervals preceding and following their aggressive behavior towards intact male mice and non-aggressive social behavior towards intact male and female mice. These data windows were not used to train the network model since they did not contain the behaviors of interest. **b)** (Left) *EN-AggINH* activity during these preceding and following intervals encoded an aggressive psychological state (red) vs. male and female non-aggressive psychological states (green; N=17 mice from the training set; F_3,67_=28.6, P<0.0001, Friedman test; post-hoc two-tailed Wilcoxon signed-rank test, P<0.005 after FDR correction). Individual animals are shown with gray lines. Data shown as mean ± s.e.m. (Right) *EN-AggINH* activity during social isolation negatively correlated with the time spent exhibiting aggressive behavior towards other males (R=-0.58; P=0.016, two-tailed Spearman’s rank correlation). Source data are provided as a Source Data file.

### EN-Aggression Inhibition (EN-AggINH) activity couples to cellular firing

*EN-AggINH* was composed of prominent theta frequency activity (4-11 Hz) in medial amygdala and beta frequency activity (14-30 Hz) in medial amygdala and prelimbic cortex (Fig. 3a-b). Prominent coherence was also observed in the theta and beta frequency bands. Indeed, when we quantified directionality across these synchronized bands via Granger causality, we saw that activity flowed from orbital frontal cortex (OFC) and primary visual cortex (V1), relayed through medial dorsal thalamus (MDThal) and then infralimbic cortex (IL), and flowed into medial amygdala (MeA) and ventral hippocampus (VHip; Fig. 3c-d, and Supplementary Fig. S5-6). Notably, these activity flow patterns involving a primary sensory region (i.e., V1) may reflect a change in encoding that occurs during aggression, or unique patterns of visual sensory input that occur during aggressive behavior.

**Figure 3.**
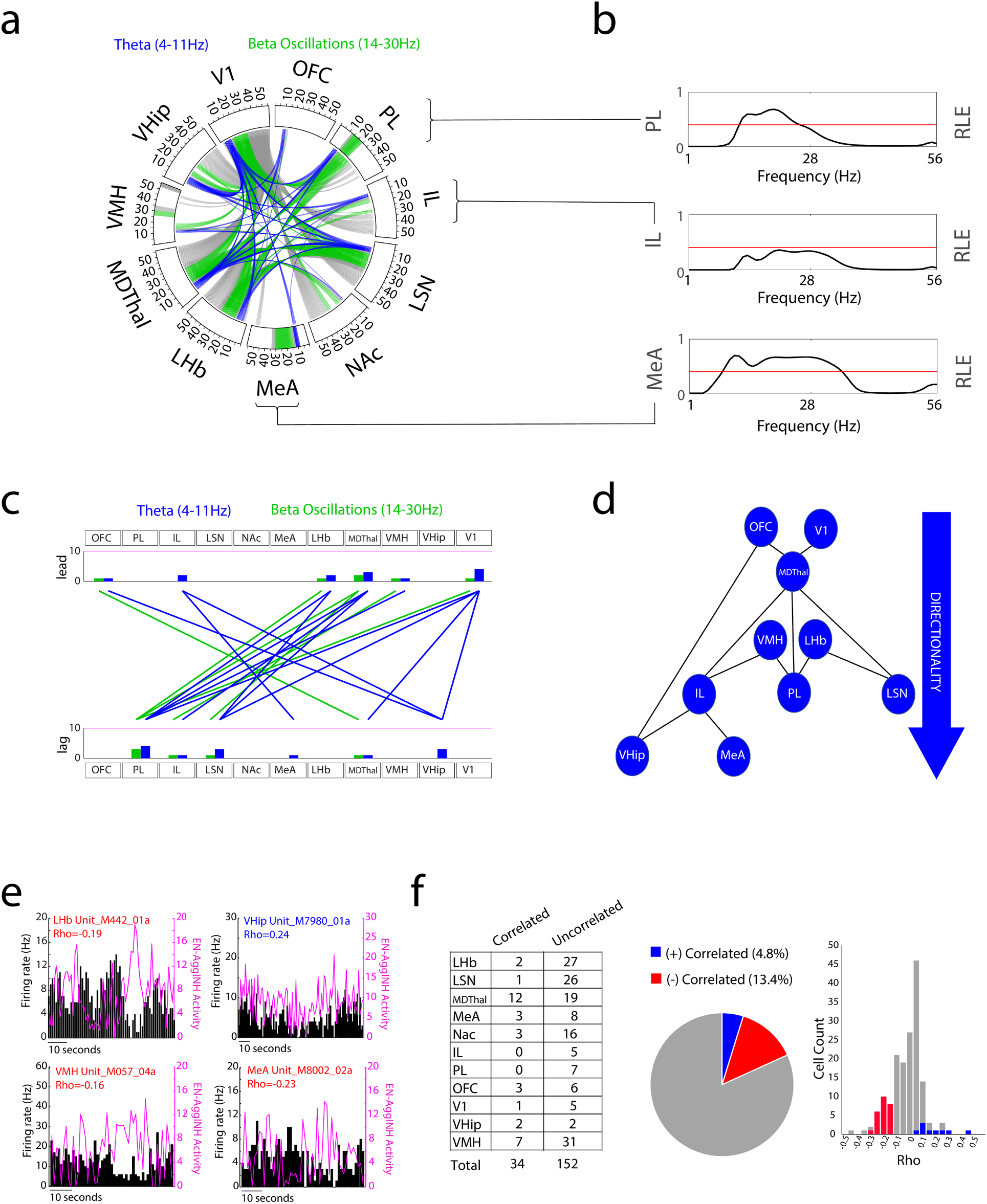
Dynamics and biological components of *EN-Aggression Inhibition (EN-AggINH)*. **a)** Prominent oscillatory frequency bands composing *EN-AggINH* are highlighted for each brain region around the rim of the circle plot. Prominent coherence measures are depicted by lines connecting brain regions through the center of the circle. The plot is shown at relative spectral energy of 0.4, corresponding to the top 15% of predictors. Theta (4-11 Hz) and beta (14-30 Hz) frequency components are highlighted in blue and green, respectively. All other frequency bands are shown in gray. **b)** Example relative LFP spectral energy (RLE) plots (predictor weights) for three brain regions corresponding to the circular plot in panel A (See Supplementary Fig. S5-6 for full description of network features). The red line corresponds to the 0.4 RLE threshold utilized for the circular plot shown in A. **c)** Directionality measures were used to depict directionality offsets (difference between A → B and B → A) within *EN-AggINH*. Prominent directionality offsets were observed across the theta and beta frequency bands (shown at spectral density threshold of 0.4 and a directionality offset of 0.3). Histograms quantify the number of leading and lagging interactions between brain regions. **d)** Schematic depicting directionality within *EN-AggINH*. **e-f)** *EN-AggINH* activity maps to cellular activity. e) Three cells from lateral habenula (LHb), ventromedial hypothalamus (VMH), and medial amygdala (MeA) showing firing activity that is negatively correlated with *EN-AggINH* activity (red) and a VHip cell showing positively correlated firing (blue). f) *EN-AggINH* activity correlated with cellular firing across the 11 recorded regions. Single- and multi-units were used for analyses. Source data are provided as a Source Data file.

Next, we verified that the activity of *EN-AggINH* truly reflects biological activity by relating the electome network to neural firing, as in previous work ^8, 42^. To achieve this, we analyzed the Spearman correlation between single and multiunit cellular activity across the implanted brain regions and the activity of *EN-AggINH*, as cell activity is an undisputed measure of biological function. We then used a permutation test to rigorously test the significance of our findings (Fig. 3e). Specifically, we shuffled cellular firing within social behavioral conditions, maintaining the relationship between cell firing and behavior. We then repeated this procedure 1000 times to generate a null distribution for which only 5% of cells would be expected to exhibit firing coupled to network activity. We found that ∼18% of cells showed firing that was coupled to the activity of *EN-AggINH*, far more than could be explained by chance (Z=4.05, p<0.0001 using two-tailed Z-test). Specifically, of the 186 cells recorded, nine (4.8%) showed firing activity that was positively correlated with *EN-AggINH* and 25 (13.4%) showed activity that was negatively correlated (Fig. 3f). Thus, most cells that showed coupling to *EN-AggINH* were inhibited when network activity increased. Notably, we also observed coupling between cellular firing and network activity when we performed high-density single-unit recordings from the anterior-medial axis of the brain using silicon probes (43 out of 473 total neurons, Z=2.41, p=0.008 using a two tailed Z-test, see Supplementary Fig. S7). Together, these analyses confirmed that *EN-AggINH* activity reflects the dynamics of cellular activity in the recorded brain regions.

### EN-Aggression Inhibition (EN-AggINH) generalizes to new biological contexts related to aggression

To further validate *EN-AggINH*, we established that activity in this network was modulated by orthogonal biological conditions that have been shown to induce or suppress aggressive behavior in mice. In most cases, we performed this analysis in new animals, which is necessary to appropriately evaluate generalization to new subjects^43^. One-tailed tests were used when prior literature provided a clear directional relationship between a given manipulation and aggressive behavior. Specifically, having established that *EN-AggINH* activity decreases during aggressive behavior, we hypothesized that causal manipulations which increased aggressive behavior would also decrease EN-AggINH activity. Similarly, we hypothesized that manipulations which decreased aggressive behavior would increase *EN-AggINH* activity.

We transformed LFP data recorded from these new sessions into *EN-AggINH* activity through our original network model. For our first validation experiment, we tested whether our network generalized to new mice on a different genetic background (Fig. 4a-b). Our approach was based on chemogenetically activating the ESR1+ cells of ventrolateral nucleus of the ventromedial hypothalamus (VMHvl), which has been shown to induce aggressive behavior towards female mice ^18, 44, 45^ – within a social context that would not normally induce aggression. We crossed CD1 strain mice with ESR1-Cre mice on a C57 strain background and expressed an excitatory DREADD (AAV-hSyn-DIO-hM3Dq) in the VMHvl of their offspring. We then treated these offspring with CNO (Clozapine N-oxide, which activates the excitatory DREADD expressed by the ESR1+ cells), and selected the subset of mice (9 out of the 13 total mice) that showed aggression toward females. Subsequently, we implanted these mice with recording electrodes to target the same brain regions used to train the network model as in our initial experiment (Fig. 4a). Following recovery, we performed behavioral and neural recordings when mice were treated with saline or CNO, in a pseudorandomized order, and then exposed to a female mouse.

**Figure 4.**
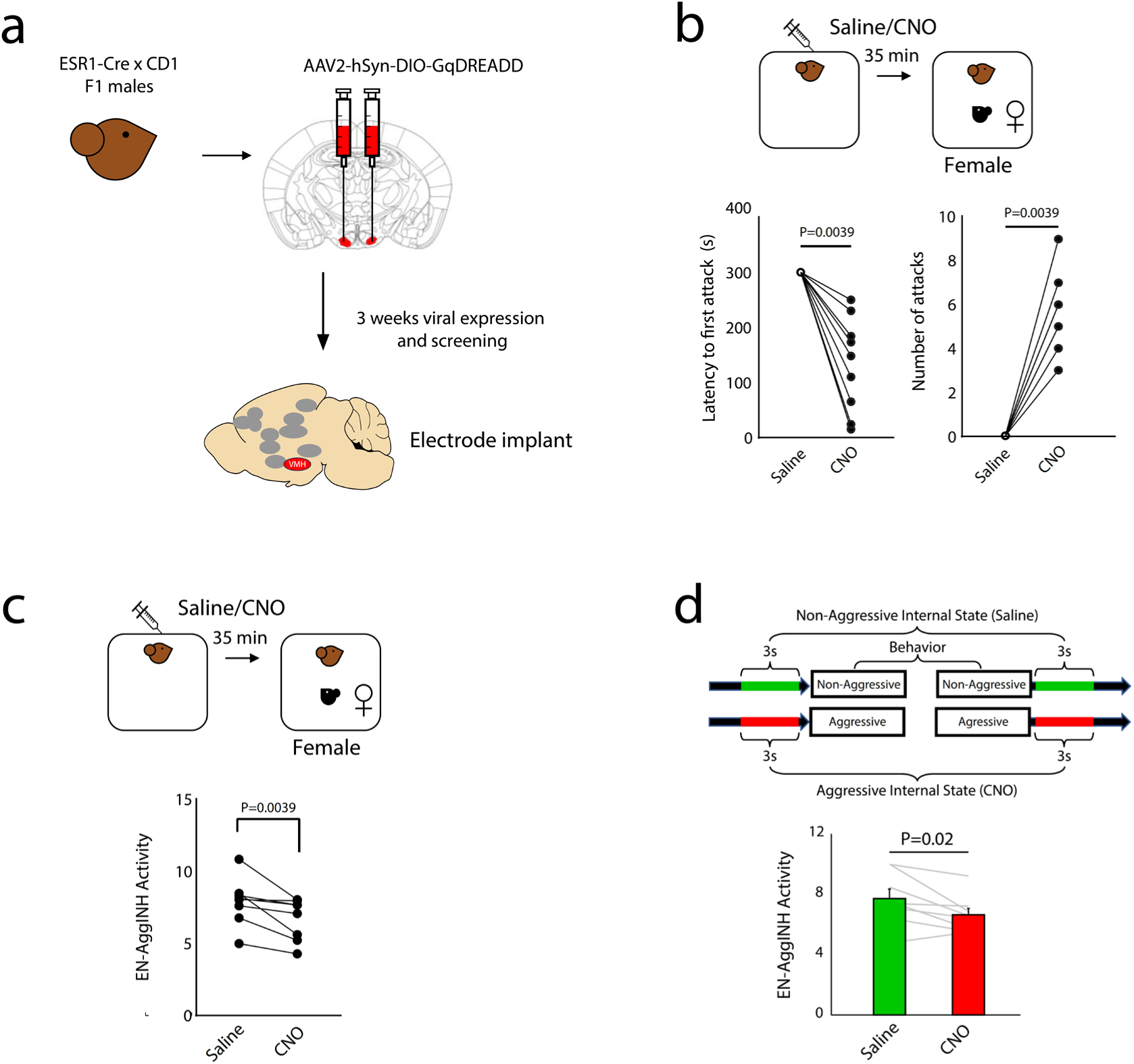
*EN-Aggression Inhibition (EN-AggINH)* encodes distinct aggression contexts. **a)** Experimental approach for causally inducing aggression via chemogenetic activation of ESR1+ cells in ventromedial hypothalamus (VMH, region in red). After three weeks for viral expression, we implanted recording electrodes to target the same brain regions used to train the network model as in our initial experiment. Following recovery, we obtained behavioral and neural recordings when mice were treated with saline or CNO. **b)** ESR1+ cell activation induced aggressive behavior towards female mice (P<0.01 using one-tailed Wilcoxon signed-rank test, N=8 mice), and **c)** decreased *EN-AggINH* activity during social interactions with female mice (P<0.01 using one-tailed Wilcoxon signed-rank test; N=8 mice) and **d)** during intervals surrounding these interactions (P<0.05 using one-tailed Wilcoxon signed-rank test). Data shown as mean ± s.e.m. Source data are provided as a Source Data file.

As anticipated, treatment with CNO induced aggressive behavior towards the female mice in the implanted mice (N=8; P=0.0039 for both attack latency and attack number using one-tailed Wilcoxon signed-rank; Fig. 4b). When we probed neural activity across the entire exposure to the female intruder, we found that treatment with CNO also suppressed *EN-AggINH* activity (N=8, P=0.0039 using one-tailed Wilcoxon signed-rank; Fig. 4c). Thus, the network model generalized to a second aggression context causally induced by a cellular manipulation, and it was robust to different genetic backgrounds. Notably, network activity was also lower during the time intervals surrounding aggressive/non-aggressive social behavior for the CNO vs. saline treatment sessions (P=0.02 using one-tailed Wilcoxon signed-rank; Fig. 4d), again demonstrating that *EN-AggINH* encoded an aggressive psychological state even while the mice were not actively engaging in aggressive behavior. Importantly, these observations also established that *EN-AggINH* does not simply encode sensory cues associated with male intruders, since the network responses observed in the CNO-treated mice were induced by a female intruder.

### EN-Aggression Inhibition (EN-AggINH) mediates aggressive behavior

We used mediation analysis to determine whether *EN-AggINH* activity putatively played a mechanistic role in suppressing aggressive behavior. Mediation analysis is a framework to determine whether the impact of a "treatment" (manipulation) on an outcome (aggressive behavior) is mediated by a change in an intermediate variable (*EN-AggINH* activity). If so, the intermediate variable is viewed, at least in part, as a mechanistic route (a mediator) for how the treatment impacts the outcome. Three components were necessary to optimally implement our mediation analysis models: a manipulation that causally modulated 1) aggressive behavior and 2) *EN-AggINH* activity, and 3) an approach to deliver the manipulation exactly when *EN-AggINH* activity would predict the emergence of aggressive behavior, i.e. at low *EN-AggINH* activity levels. We chose to build such an approach based on optogenetic stimulation of mPFC, since we had previously found that such a manipulation causally suppressed aggressive behavior ^22^.

First, we built a closed-loop system that estimated the activity of *EN-AggINH* in real time (Fig. 5a). This approach employed a new network encoded solely based on power and coherence measures (i.e., a reduced network, Fig. 5b), because the processing time to calculate Granger causality was prohibitive for real-time implementations. While this new network lacked the full interpretive power of *EN-AggINH*, it enabled us to predict aggressive behavior in real time to a similar degree as *EN-AggINH* did (N=9; P=0.43 using two-tailed paired Wilcoxon signed-rank; Fig. 5c). In principle, when the activity of *EN-AggINH* fell below an established threshold (signaling the onset of aggressive behavior, see methods section: Optogenetic stimulation), our closed-loop approach would deliver a one-second light stimulation (5mW, 20Hz, 3ms pulse width) to mPFC. To verify that this real-time estimation system worked as designed, we tested whether light stimulation corresponded with a decrease in *EN-AggINH* activity (i.e., the full network). Indeed, network activity was significantly lower one second prior to stimulation than it was two seconds prior to stimulation, demonstrating that our approach using the reduced encoder successfully identified when the *EN-AggINH* activity decreased below the threshold that signaled the onset of aggression (N=9; P<0.005 for within-subject comparison of *EN-AggINH* activity 1 vs. 2 seconds prior to yellow light stimulation using one-tailed Wilcoxon signed-rank test, Fig. 5d). Importantly, we found that mPFC stimulation acutely increased *EN-AggINH* activity (N=9, P<0.01 for comparison of *EN-AggINH* activity one second after blue vs. yellow stimulation, using one-tailed Wilcoxon signed-rank test, see Fig. 5d). Thus, our closed-loop stimulation approach satisfied two of the components needed to implement our mediation approach.

**Figure 5.**
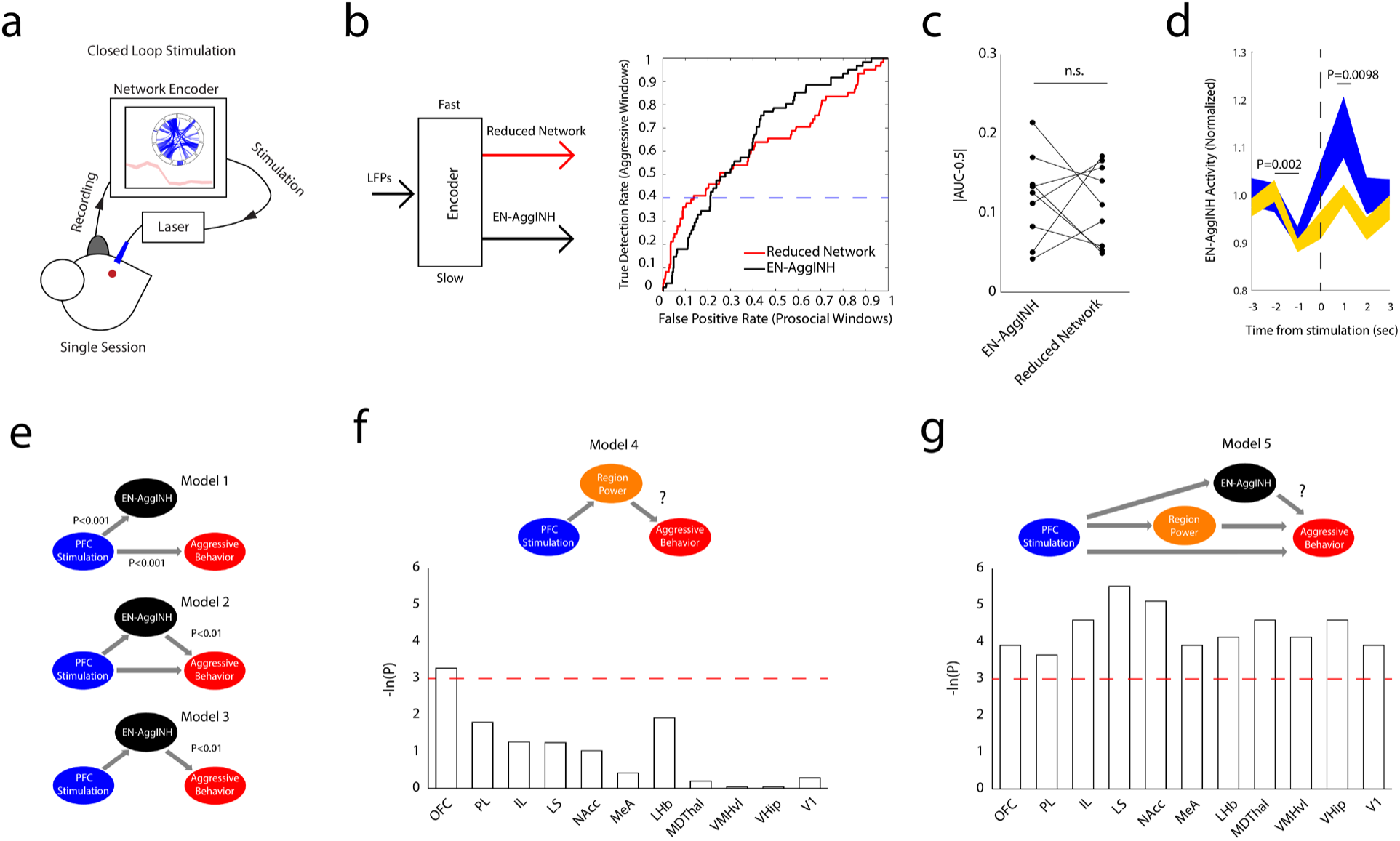
*EN-Aggression Inhibition (EN-AggINH)* activity is causally related to aggression. **a)** Schematic for closed-loop manipulation of *EN-AggINH* activity. **b)** Real-time estimation of aggression. Receiver operating characteristic depicting detection of aggressive behavior in a mouse using *EN-AggINH* activity vs. real-time reduced encoder (i.e., based solely on power and coherence features) is shown to the right. Dashed blue line corresponds to the established detection threshold. **c)** Detection of aggression using reduced encoder vs. *EN-AggINH* across mice (N=9 mice; P=0.43 using two-tailed paired Wilcoxon signed-rank). **d)** *EN-AggINH* activity in response to blue and yellow light stimulation during closed-loop manipulation. Network activity significantly decreased one second prior to yellow light stimulation (N=9, P<0.005 using one-tailed Wilcoxon signed-rank test for within-subject comparison of *EN-AggINH* activity 1 vs. 2 seconds prior to yellow light stimulation; note that activity was normalized to the average network activity during isolation for each mouse). Activity was also higher one second after stimulation with blue light vs. yellow light (P<0.01 using one-tailed Wilcoxon signed-rank test). **e)** Directed graph with the inferred modes of action derived from mediation analysis. Model 1: There is a causal relationship from stimulation to behavior and from stimulation to *EN-AggINH* expression (P<0.001 using Wilcoxon one-tailed signed-rank test for network activity, and two-tailed and paired t-tests for behavior). Model 2: *EN-AggINH* is a mediator from stimulation to behavior (P<0.01 using two-tailed nested logistic regression models, likelihood ratio test). Model 3: *EN-AggINH* activation is the primary mechanism whereby mPFC stimulation suppresses aggression (P<0.01 using two-tailed average causal mediation effect). **f-g)** Directed graphs testing f) theta power within single regions as the primary mechanism whereby mPFC stimulation suppresses aggression (model 4) and g) *EN-AggINH* activation as the primary mechanism whereby mPFC stimulation suppresses aggression when single-region power is included as an intermediary (model 5). The uncorrected P values for each brain area in both models are shown below as -ln(P), where the red dashed line corresponds to P=0.05. Source data are provided as a Source Data file.

For the final component, we tested whether increasing *EN-AggINH* activity via mPFC stimulation as the brain transitioned into a putative aggressive psychological state would suppress aggressive behavior. Our closed-loop approach preferentially induced optogenetic stimulation during aggressive interactions, compared to non-aggressive social interaction with females, establishing its selectivity (P = 0.002 using one-tailed Wilcoxon signed rank test; Fig. 6a, left). Importantly, closed-loop stimulation significantly suppressed aggressive behavior (N=9 mice that were not used to train the initial model; t8=6.1, P=0.0003, comparing blue vs. yellow light stimulation using two-tailed paired t-test for aggressive behavior, significance determined by FDR correction; Fig. 6a, right;). Thus, our closed-loop manipulation suppressed aggressive behavior, satisfying the remaining component needed to implement our mediation analysis approach.

**Figure 6.**
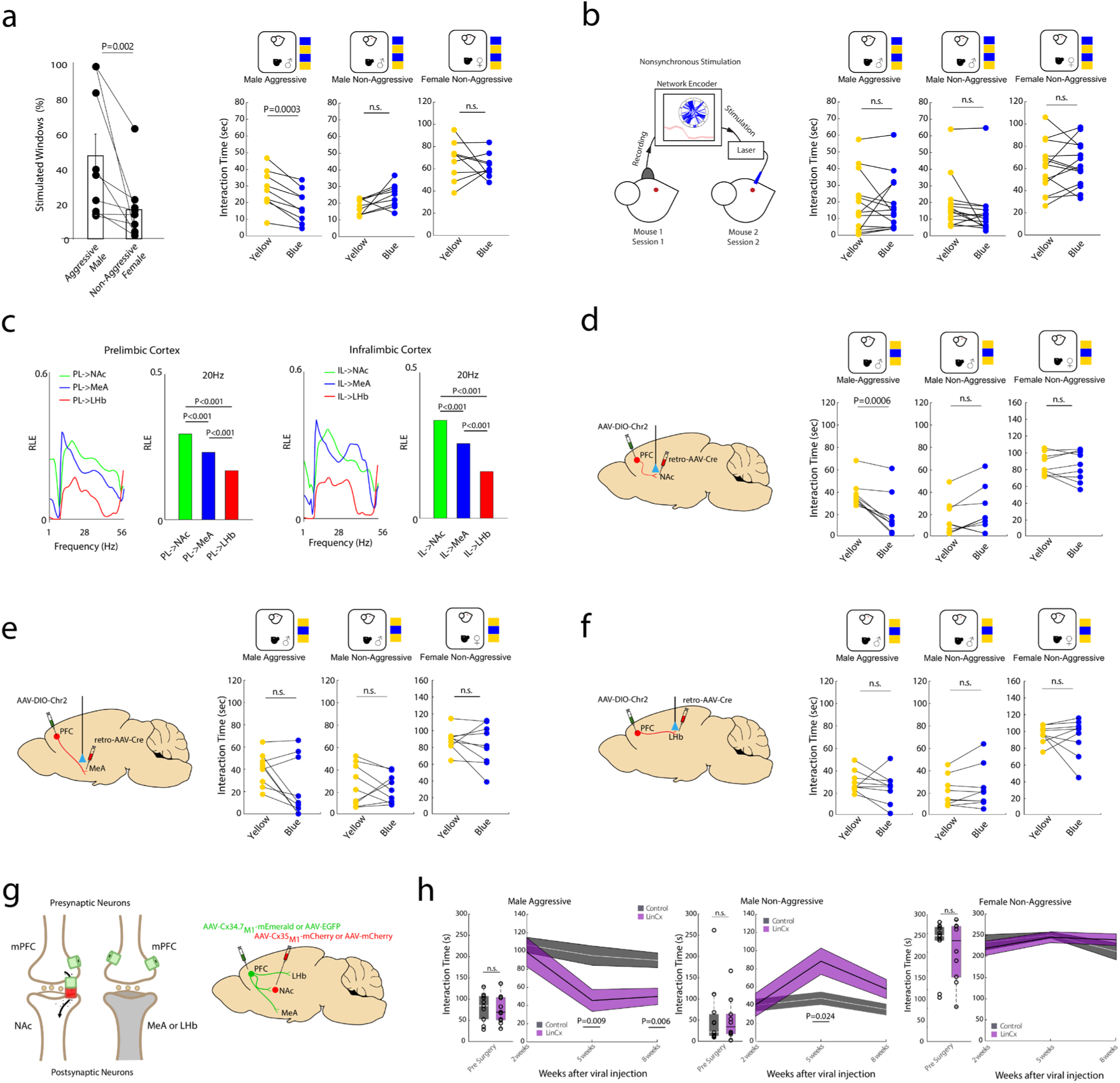
Validation of spatiotemporal features of *EN-Aggression Inhibition (EN-AggINH)*. **a)** Percentage of windows stimulated during social behaviors using a closed-loop approach, which preferentially induced optogenetic stimulation during aggressive interactions, compared to non-aggressive social interaction with females (P=0.002 using one-tailed Wilcoxon signed-rank test, left). Data shown as mean ± s.e.m. Behavioral effects of closed-loop stimulation are shown on the right (blue and yellow light stimulation compared using two-tailed paired t-test, significance determined using FDR correction; N=9 mice). **b)** Schematic for nonsynchronous control stimulation (left). Nonsynchronous stimulation does not impact aggressive or non-aggressive social behavior towards males or females (N=14 mice). **c)** Directionality for mPFC-dependent subcircuits within *EN-AggINH* (shown as relative LFP spectral energy or RLE, see also Supplementary Fig. S6). P<0.01 using bootstrapping procedure over 1000 repeats. **d)** Viral targeting strategy (left) and behavioral impact of PL → NAc circuit stimulation (right; N=9 mice). **e)** Viral targeting strategy (left) and behavioral impact of PL → MeA circuit stimulation (right; N=9 mice). **f)** Viral targeting strategy (left) and behavioral impact of PL → LH circuit stimulation (right; N=9 mice). For all panels B, D-F: P<0.01 using two-tailed paired t-test. Order of blue and yellow light stimulation trials is shown next to the right of each social condition diagram in panels A-B, D-F. **g)** Left: Schematic of electrical synapse formation for circuit-selective LinCx editing. Cx34.7_M1_ (green hemichannel) and Cx35_M1_ (red hemichannel) show heterotypic docking with each other, but not homotypic docking between two Cx34.7_M1_ hemichannels. The synapse formed between them rectifies in the Cx34.7_M1_ to Cx35_M1_ direction. Right: Schematic of Cx34.7_M1_ and Cx35_M1_ vs. control fluorophore targeting the mPFC and NAc specifically. **h)** Impact of LinCx editing the mPFC-NAc circuit on aggressive (left) and non-aggressive behavior towards intact males (middle), and non-aggressive behavior towards females (right). Data is plotted as mean ± s.e.m. at 2, 5, and 8 weeks after viral injection (P<0.05 using repeated measures ANOVA followed by two-tailed unpaired t-test across groups; N=10 LinCx-edited mice and N=11 control mice). Box and whisker plots to the left of each plot show social behavior in CD1 animals following group assignment, but prior to viral injection. No differences were observed across groups for any of the three social behaviors (P>0.05 for all comparisons across groups, unpaired two-tailed t-test). For the box and whisker plots, the central mark is the median, the edges of the box are the 25th and 75th percentiles, the whiskers extend to the most extreme data points the algorithm does not consider to be outliers. Individual data points are shown as circles. Source data are provided as a Source Data file.

We first used the classic Baron and Kenny approach ^46^ to determine whether *EN-AggINH* activity mediates the effect of neurostimulation on aggressive behavior. According to this statistical approach, there is a mediated effect of network activity on behavior if three conditions are met: 1) stimulation modulates network activity, 2) network activity correlates with behavior, and 3) modeling the behavior from network activity and stimulation together is better than modeling behavior from stimulation alone. Indeed, we had identified a significant direct effect of stimulation on aggressive behavior (see Fig. 6a, right) and network activity (see Fig. 5d). To optimally match the analytical conditions between the treatment and control cases for subsequent analysis, we used windows during the closed-loop stimulation procedure where the laser was triggered and then compared blue laser stimulation (treatment) to yellow laser stimulation (control). Thus, the data points used for our mediation analysis predicted imminent or ongoing aggressive behavior, and network activity prior to the stimulation in both the control (yellow light) and treatment (blue light) case were similar. A statistical model of behavior using network activity and stimulation (see Fig. 5e, model 2) significantly outperformed the model using only stimulation (see Fig. 5e, model 1; nested logistic regression models, P<0.01, likelihood ratio test), satisfying the necessary conditions to show that *EN-AggINH* is a mediator.

After establishing that *EN-AggINH* activity mediated the impact of mPFC stimulation on behavior, we set out to evaluate the significance of the average causal mediation effect (ACME) and the average direct effect (ADE) during the same stimulated closed-loop windows using causal mediation analysis ^47^. ACME is the causal effect of stimulation on behavior due to the change in *EN-AggINH* activity (see Fig. 5e, model 3), and ADE is the causal effect on behavior from mPFC stimulation not explained by the change in *EN-AggINH* activity. We found that there was a significant ACME (P<0.01), but not a significant ADE (P=0.48). This analysis suggested that *EN-AggINH* activation is the primary mechanism whereby mPFC stimulation suppresses aggression.

Next, we tested different mediation pathways to determine whether the suppressive effects of stimulation on aggression are specifically mediated by *EN-AggINH* rather than alternative neural processes. First, we evaluated whether theta power in any of the 11 different brain regions could serve as a mediator in lieu of *EN-AggINH* activity (Fig. 5f, model 4). We chose this frequency band since it was prominently featured in *EN-AggINH* and also in the electome network we previously found to encode social appetitive behavior^8^. Across these 11 single-region models, only OFC had a significant average causal mediation effect (uncorrected p-value of 0.038, see Fig. 5f), but this model did not survive a correction for multiple comparisons. Additionally, its ACME estimate was dwarfed by the size of the ACME estimate for *EN-AggINH* (the mean estimate for the *EN-AggINH* model was 49.7% larger). This evidence suggests that *EN-AggINH* is a much better mediator than any of these potential single-region ’biomarkers’ by themselves.

After failing to identify any significant mediation effect of theta activity within each of the eleven brain regions, we tested whether including theta activity as an intermediary in our causal graph would disrupt *EN-AggINH’s* role as a mediator in aggressive behavior (Fig. 5g, model 5). Here, we corrected for the role of theta power in the model of how *EN-AggINH* changes as a function of stimulation, as well as correcting for theta power in forecasting aggressive behavior. As such, this framework dictates that *EN-AggINH* cannot mediate behavior that is already explained by changes in theta power in a specified region. When we ran eleven models, one model for each brain region, we found that *EN-AggINH* still significantly mediated aggressive behavior in all of them (P<0.05 for all models; see Fig. 5g, bottom). Thus, even after accounting for these potential intermediate variables, our findings still supported *EN-AggINH* as a mediator of aggressive behavior. Consistent with this finding, our mediation analysis performed using data from the ESR1-Cre experiment showed that *EN-AggINH* also mediated the impact of CNO treatment (see Supplementary Fig. S8). Thus, *EN-AggINH* reflects the internal brain state that suppresses aggressive behavior.

### Validation of temporal activity and spatial spectral features of EN-AggINH

We validated the temporal activity and spatial spectral features of *EN-AggINH* by establishing that they could be utilized to selectively suppress aggression. Specifically, after determining that our closed-loop manipulation suppressed aggression, we also quantified the impact of this stimulation protocol on non-aggressive interactions with other male and female mice. We found that closed-loop mPFC stimulation had no impact on non-aggressive behavior towards the intact C57 males (N=9; t8=-2.3, uncorrected P=0.049 comparing blue vs. yellow light stimulation using two-tailed paired t-test for non-aggressive behavior, significance determined by FDR correction, Fig. 6a, right, 2^nd^ graph). No stimulation-induced differences in non-aggressive behavior were observed during exposure to female mice (t_8_=0.74, P=0.48 using two-tailed paired t-test, significance determined by FDR correction, Fig. 6a, right, 3^rd^ graph). Interestingly, while the open-loop protocol induced hyperactivity during open field exploration (Supplementary Fig. S9a, left), no differences in gross locomotor behavior were observed during closed-loop mPFC stimulation (Supplementary Fig. S9a, right). Thus, closed-loop mPFC stimulation selectively reduced aggression without affecting other social behaviors or gross locomotor behavior.

Next, we verified that this selective modulation of aggression was due to synchronization of light stimulation with endogenous *EN-AggINH* activity, and not simply due to the dynamic pattern of stimulation delivered using this method. We performed an additional control experiment where mPFC stimulation patterns in new animals were yoked to network activity recorded from another mouse’s brain, analogous to a ’sham’ in neurofeedback experiments (i.e., nonsynchronous stimulation; Fig. 6b). Nonsynchronous stimulation failed to suppress aggressive behavior (F_1,21_=4.87, P =0.039 for light type × stimulation pattern effect for exploratory analysis using a mixed effects, two-way ANOVA; t_13_=0.09, P=0.93 for nonsynchronous stimulation effects on aggressive behavior towards intruder male mice using post-hoc two-tailed paired t-test, significance determined by FDR correction; see Fig. 6b, right, 1^st^ graph), verifying that the suppression of aggressive behavior driven by closed-loop stimulation was indeed due to delivery of stimulation timed to endogenous *EN-AggINH* activity. Incidentally, nonsynchronous stimulation had no impact on non-aggressive social behavior towards intact males or females (N=14; t_13_=1.79, P=0.097; and t_13_=0.54, P=0.60, for interaction with males and females, respectively, comparing blue vs. yellow light stimulation using two-tailed post-hoc paired t-tests, significance determined by FDR correction; see Fig. 6b, right, 2^nd^ and 3^rd^ graphs). Thus, we validated the temporal activity component of *EN-AggINH*.

After establishing that we could selectively reduce aggression by temporally targeting mPFC based on the activity level of *EN-AggINH*, we tested whether we could reduce aggression by spatially targeting stimulation based on the sub-components of mPFC output circuity that composed the network. Our goal was to identify sub-components of the *EN-AggINH* whose information flow changes most as the system transitions between aggression states to find promising targets for control. Our initial visualization of *EN-AggINH* was constrained to the absolute information flow at the strongest synchronies (Fig. 3c-d). On the other hand, relative LFP spectral energy (RLE) is the weight of each directionality feature of *EN-AggINH* normalized to the sum of the weight of that feature across all the learned electome networks. As such, the directionality RLEs quantify the magnitude to which a specific circuit increases its output as *EN-AggINH* activity increases (Supplementary Fig. S6). We therefore identified target circuits by looking at the RLE of the *EN-AggINH* directionality features that emerged from prelimbic (PL) and infralimbic cortex (IL) cortex. Because *EN-AggINH* activity decreases during aggression, a high directionality RLE also corresponds with the circuits within *EN-AggINH* that show the strongest reduction in information flow when aggression emerges.

We focused our analysis on the directionality between mPFC [prelimbic (PL) and infralimbic cortex (IL)] to nucleus accumbens (mPFC → NAc), medial amygdala (mPFC → MeA) and lateral habenula (mPFC → LHb), since these circuits consisted of monosynaptic projections, enabling direct targeting using optogenetics. We then quantified the 20Hz RLE of these mPFC output circuits (Fig. 6c), since increasing mPFC activity at this frequency via optogenetic stimulation was sufficient to suppress aggressive behavior (Fig. 1a) and increase *EN-AggINH* activity (Fig. 5d). *EN-AggINH’s* RLE was highest for mPFC → NAc and then mPFC → MeA circuitry and lowest in the mPFC → LHb circuit. Thus, a prominent decrease in information flow in mPFC → NAc and mPFC → MeA circuitry was associated with aggression, while no such change was observed in mPFC-LHb activity [P<0.001 for pairwise comparison of relative LFP spectral energy across these six circuits (analysis performed separately for PL- and IL-dependent circuitry] using bootstrapping, see methods)]. Given this observation, we reasoned that driving mPFC → NAc or mPFC → MeA activity at 20Hz should selectively suppress aggression, while driving mPFC → LHb activity should not.

We causally activated these three circuits at 20Hz and measured their impact on social behaviors. To selectively stimulate the terminals of mPFC neurons in each target region (NAc, MeA, or LHb), we injected mice with a retrograde AAV-Cre (rAAV-Cre) virus in one target region and an AAV-DIO-channelrhodopsin-2 virus in mPFC. A stimulating fiber was placed above the target region injected with rAAV-Cre. Social behavior was quantified during 20Hz stimulation with yellow vs. blue light (5mW, 20Hz, 3ms pulse width). Blue light stimulation of mPFC → NAc decreased aggression toward male C57 mice (t8=5.5, P=0.0006 for blue vs. yellow light using two-tailed paired t-test; N=9 mice, significance determined by FDR correction), but had no impact on non-aggressive social behavior towards the male C57 mice (t_8_=2.01, P=0.08 using a two-tailed paired t-test) or social behavior towards female C57 mice (t_8_=0.39, P=0.71 using a two-tailed paired t-test; see Fig. 6d, right). Though mPFC → MeA tended to decrease aggression toward male C57 mice, these differences did not reach statistical significance (t_8_=2.24, P=0.056 for blue vs. yellow light using a two-tailed paired t-test; N=9 mice, significance determined by FDR correction; see Fig. 6e, right, 1^st^ graph). Again, no difference in non-aggressive social behavior was observed towards male C57 mice (t_8_=0.58, P=0.58 using a two-tailed paired t-test) or females (t_8_=0.80, P=0.45 using a two-tailed paired t-test; see Fig. 6e, right). Finally, blue light stimulation of mPFC → LHb stimulation had no impact on aggression toward male C57 mice (t_8_=1.05, P=0.32; using two-tailed paired t-test, N=9 mice), non-aggressive social behavior towards C57 males (t_8_=1.42, P=0.19 using two-tailed paired t-test), or social interaction with C57 females (t_8_=0.45, P=0.67 using two-tailed paired t-test; see Fig. 6f, right). These results demonstrated that directly stimulating the mPFC subcircuit which showed the greatest decreases in aggression-related activity (i.e., mPFC → NAc) causally suppressed aggression. On the other hand, stimulating a mPFC subcircuit with minimal activity changes during aggression (i.e., mPFC → LHb) had no impact on social behavior towards male mice. Together, these findings validated the spatial spectral features of *EN-AggINH*.

Having established that acute stimulation of the mPFC → NAc circuit selectively diminished aggression, we tested whether editing this circuit using LinCx chronically suppressed aggressive behavior. LinCx is composed of two gap junction hemichannels, Cx34.7_M1_ and Cx35_M1_, which exclusively dock with each other^48^. Each of the hemichannels can be targeted to distinct cell membranes that are in close proximity to yield an electrical synapse between the pair of cells (Fig. 6g, left), increasing the coupling of neural activity between them^48^. After screening, we assigned aggressive male CD1 mice to experimental and control groups in a manner that balanced social behavior (aggressive and non-aggressive) between groups. After another presurgical behavioral screening, experimental mice were injected with AAV-CaMKII-Cx34.7_M1_-2xHA-T2A-mEmerald in mPFC and AAV-CaMKII-Cx35_M1_-FLAG-T2A-mCherry in NAc. Control mice were injected with AAV-CaMKII-GFP in mPFC and AAV-CaMKII-mCherry in NAc (Fig. 6g, right). We then quantified aggressive and non-aggressive social behavior serially at 2-, 5-, and 8-weeks post-injection (Fig. 6h). Notably, strong trafficking of Cx34.7_M1_ from mPFC to NAc (and thus LinCx editing) is not observed until more than 3 weeks after the initial surgery^48^. Importantly, the second pre-surgical screening of aggressive and non-aggressive behavior towards intact males and females found no difference between the experimental and control groups (t_19_=0.35, P=0.73 for group comparison of aggressive behavior towards intact males, t_19_=0.23, P=0.82 for group comparison of non-aggressive social behavior towards intact males, and t_19_=0.57, P=0.57 for group comparison of non-aggressive social behavior towards females, all using a two-tailed t-test; N=10 LinCx-edited mice and N=11 control mice), demonstrating the robustness of our group assignments (see Fig. 6h). Our analysis using a three-way ANOVA found a significant group x social condition effect on social interaction time (F_4,168_=2.097 and P=0.08 for group × social condition × week effect using mixed-effects model three-way ANOVA; F_2,168_=7.38 and P=0.0008 for group × social condition effect using mixed-effects model three-way ANOVA). Indeed, editing the mPFC → NAc circuit using LinCx decreased aggressive behavior towards intact males (F_2,38_=2.85 and P=0.07 for group × week effect using mixed-effects model ANOVA; F_1,19_=6.82 and P=0.017 for group effect, F_2,38_=8.6 and P<0.001 for week effect; significance determined by a Benjamini-Hochberg false discovery rate (FDR) correction; T_19_=2.92 and P=0.009; T_19_=3.13 and P=0.006 for post-hoc comparisons across groups at 5 and 8 weeks after injection, respectively, using an unpaired two-tailed t-test; N=10 LinCx-edited mice and N=11 control mice; Fig. 6h, left). These mice showed increased non-aggressive social behavior towards intact male intruders at 5, but not 8 weeks after injection (F_2,38_=5.26 and P=0.01 for group × week effect using mixed effects model ANOVA; significance determined by a Benjamini-Hochberg false discovery rate (FDR) correction; T_19_=2.46 and P=0.024; T_19_=1.79 and P=0.090 for post-hoc comparisons across groups at 5 and 8 weeks after injection, respectively, using an unpaired two-tailed t-test; Fig. 6h, middle). Finally, we observed no difference in social behavior towards females (F_1,19_=0.02 and P=0.88 for group effect; F_2,38_=2.05 and P=0.14 for group × week effect using mixed effects model ANOVA; significance determined by a Benjamini-Hochberg false discovery rate (FDR) correction; Fig. 6h, right). Consistent with our closed-loop mPFC stimulation findings, LinCx editing did not induce hyperactivity during open field exploration (Supplementary Fig. S9b). Taken together, these findings showed that editing the mPFC → NAc chronically suppressed aggressive behavior, while leaving non-aggressive social behavior intact.

## DISCUSSION

We set out to identify a neural circuit that could be targeted to selectively decrease aggressive behavior while leaving pro-social behavior intact. To achieve this outcome, we investigated the neural mechanisms that underlie when a mouse engages in aggressive vs. pro-social behavior. We reasoned that aggressive behaviors emerge when the brain is in an aggressive psychological state, and that pro-social behaviors emerge when it is not. Thus, we used machine learning to discover the mesoscale neural architecture of the brain when an animal expressed aggressive behavior versus when it expressed non-aggressive social behaviors. The network we identified incorporated distributed neural dynamics spanning all recorded regions. Distributed neural dynamics are also observed in well-defined internal brain states such as sleep, which are characterized by coordinated activity changes across widespread brain regions^49, 50^. For multiple regions, differences were observed in local oscillatory power, while others exhibited differences in coherence between a broader collection of regions. No brain region showed prominent activity contributions across all frequencies. Rather, each brain region showed selectivity in the frequencies of oscillations that contributed to the network. For example, prelimbic cortex showed strong activity in the beta frequency range, while medial amygdala showed strong activity in the beta and theta frequency ranges. Most importantly, the brain state that was identified across multiple contexts associated with aggression was better captured by an integrated network composed of activity recorded across 11 brain regions, rather than the independent activity within each brain region.

Interestingly, this aggressive brain state was encoded by decreased activity in the network. Given that we identified more cells that increased their firing rates as network activity decreased, the discovery of a network that decreases its activity during aggression does not indicate that overall brain activity is suppressed during aggressive states. Rather, these findings argue that the aggressive state is encoded by a network that decreases its activity relative to when mice are socially isolated or engaged in pro-social behavior (Fig. 4B, left). Indeed, our data suggested that several common regions/circuits were activated during aggressive and pro-social behavior. These common regions/circuits need not be reflected in our network since our model was trained to differentiate aggressive from non-aggressive social behavior. Nevertheless, our discovery of a network that decreased its activity during aggression raises the intriguing hypothesis that the brain actively inhibits aggression during pro-social engagement. When activity in this inhibition network is suppressed, aggressive behavior emerges.

This interpretation is supported by our validation experiments where we chemogenetically activated ESR1+ neurons in ventromedial hypothalamus, which induced the aggressive brain state (i.e., suppressed *EN-AggINH* activity). When CNO-treated mice were then exposed to a stimulus that would generally induce non-aggressive social behavior (i.e., a female mouse), aggressive behavior emerged. Thus, the presence of the aggressive brain state changed the mapping between sensory input and behavior output. Similarly, closed-loop optogenetic stimulation of mPFC biased mice towards exhibiting non-aggressive social behavior when they were exposed to a stimulus that would generally induce aggressive behavior (i.e., a male intruder), as well as decreased the aggressive brain state (i.e., increased *EN-AggINH* network activity). Finally, our mediation analyses suggest that the brain state represented by *EN-AggINH* mediates the suppressive effects of mPFC stimulation on aggressive behavior.

Though behavioral output has been classically utilized to infer the internal state of a brain, we reasoned that an internal aggressive psychological state was also likely present during intervals immediately preceding and following aggressive behavior. Thus, we tested whether our network showed distinct activity profiles in the time intervals surrounding aggressive behaviors. Indeed, network activity was lower during intervals surrounding aggressive behavior compared to intervals surrounding non-aggressive behavior. Strikingly, we also found that network activity when animals were isolated in their home cage encoded their trait aggression. Thus, the network did not simply encode aggressive action since its activity was observed separately from behavior. Rather, the network encoded an internal aggressive psychological state.

Here, we framed internal brain states as a collection of functions that transform sensory input into behavior. Indeed, we found that when *EN-AggINH* activity is suppressed, the brain transforms both male and female social sensory cues into aggressive behavior. It is also widely appreciated that internal and external sensory input can also cause the brain to transition from one internal state to another. Along this line, we found that exposure to male mice could promote an aggressive psychological internal state in CD1 mice even prior to aggressive behavior, while exposure to a female mouse did not (under normal conditions). In this framework, one would also anticipate that modulation approaches that decrease aggressive behavior could mediate their effect by driving the brain out of the aggressive state, which is represented by low *EN-AggINH* activity.

The ability to modulate aggressive behavior in this way could present a breakthrough for the treatment of multiple neuropsychiatric disorders – including mood disorders, psychotic disorders, neurodevelopmental disorders, and neurodegenerative disorders – with deficits in regulating social behavior such as aggression. While pharmacological approaches have been implemented to suppress aggressive behavior towards self and others, many of these strategies act by sedating the individual and can disrupt aspects of pro-social function. Our discovery of a brain network that encoded an aggressive state raises the potential for novel approaches to suppress aggressive behavior that spare pro-social behavior. Indeed, compared to a standard open-loop stimulation protocol which suppressed both aggressive behavior toward intact males and non-aggressive pro-social behavior toward females (Fig. 1a), our closed-loop stimulation approach spared non-aggressive social behavior towards females. Intriguingly, like other open-loop mPFC stimulation studies ^51, 52^, our 20Hz stimulation protocol induced behavioral hyperactivity in experimental mice (Supplementary Fig. S9a, left). On the other hand, our closed-loop stimulation protocol did not (Supplementary Fig. S9a, right). Thus, our findings also show that closed-loop stimulation may limit off-target behavioral effects that are induced by classic stimulation approaches.

Our closed-loop stimulation approach was developed using a neural network-based approximation technique for which the features were substantially constrained relative to dCSFA-NMF. Nevertheless, we found that the reduced encoder was sufficient to identify the precise time windows when the brain transitioned into an aggressive psychological state, as marked by a decrease in *EN-AggINH* activity. In the future, novel approaches may allow for further improvement in the precision of our real-time stimulation approach, including fast approximation by surrogate neural networks. Such an approach may enable the implementation of multiple real-time networks that predict both aggressive and pro-social states concurrently to actuate our closed-loop system and further suppress aggressive behavior relative to pro-social behavior. Interestingly, our current findings also pointed to a network that exhibits increased activity during aggressive behavior (electome network #6, see Fig. 1f and Supplementary Fig. S10). Though the network failed to encode the urine paradigm and thus did not generalize across aggressive contexts, it is possible that it contains activity that synergizes with *EN-AggINH* to encode aggressive social states more optimally. If future studies demonstrate this network’s potential, a reduced encoder for electome network #6 could also be integrated to further optimize our closed-loop approach to selectively suppress aggression.

Extending our mPFC stimulation studies, we used the spatiotemporal architecture of *EN-AggINH* to identify an mPFC output circuit to NAc that was prominently suppressed during aggression. Activating this mPFC → NAc circuit at 20Hz (open-loop optogenetic stimulation) reduced aggression while sparing pro-social behavior towards intact males and females. Stimulating another mPFC output circuit to LHb that was relatively less suppressed in *EN-AggINH* failed to reduce aggression. Having identified this circuit specificity, we utilized LinCx to edit the mPFC → NAc circuit, which chronically suppressed aggression and temporarily increased pro-social behavior towards intact males. Additionally, this manipulation spared pro-social behavior towards female mice. Thus, by mapping the distributed brain network that encodes aggressive behavior and a generalized aggressive psychological state, performing multiple causal manipulation experiments to validate the spatiotemporal features of this network, and deploying targeted circuit editing using LinCx, we established a neural circuit that could be targeted to diminish aggressive behavior while leaving pro-social behavior intact.

Overall, our findings establish a generalized network-level signature whereby the brain suppresses aggression via active inhibition. Moreover, they highlight the exciting therapeutic potential for state-specific (i.e., closed-loop) neuromodulation to regulate internal states and for neural circuit editing to chronically and selectively suppress pathological social behavior.

## MATERIALS AND METHODS

### Animal care and use

All procedures were approved by the Duke University Institutional Animal Care and Use Committee in compliance with National Institute of Health (NIH) Guidelines for the Care and Use of Laboratory Animals. Mice were maintained on a reverse 12-hr light cycle with *ad libitum* access to food and water.

45 six-month-old, retired breeder male CD1 strain mice (Charles River Laboratories, Wilmington, Massachusetts) were used to discover a network that encoded aggression, hereafter called *EN-AggINH*. Another 226 male CD1 mice were used to examine behavioral and network responses to optogenetic stimulation. Mice were singly housed with enrichment.

ESR1-Cre male mice on a C57BL/6J background were provided by Scott Russo for the chemogenetic experiments. These mice were crossed with female CD1 strain mice (Charles River Laboratories, Wilmington, Massachusetts) in the Bryan Vivarium at Duke University to obtain F1 offspring. 13 fourteen-week-old virgin ESR1-Cre+/CD1 male offspring were used to validate *EN-AggINH*. All F1 offspring were group-housed 2-5 mice per cage until they received viral injections in the ventromedial hypothalamus at 7-8 weeks. After surgery, these mice were singly housed with enrichment.

60 six-month-old, retired breeder male CD1 strain mice were used for the LinCx editing experiment. Mice were singly housed with enrichment

All intruder mice (C57BL/6J: 2-7 intact males, 2-7 females, and 2-7 castrated males per experimental mouse) were 7-14 weeks old. These mice were purchased from Jackson Laboratories (Bar Harbor, Maine) and were housed 5 per cage with enrichment. All behavioral testing and neurophysiological recordings occurred during the dark cycle.

### Castration of C57BL/6J male mice

18 male mice were anesthetized with 1% isoflurane. The scrotal sac was sanitized with betadine and 70% ethanol. The testes were then moved into the sac by gently palpating the lower abdomen. Next, an incision was made in the sac and the testes were extracted. After blood flow was cut off to the testes using a thread tourniquet, the testes were severed from the mouse. The remaining fatty tissue was placed back into the scrotum, which was then sutured. Mice were allowed 10 days for recovery prior to experimental use.

### Electrode implantation surgery

Mice were anesthetized with 1% isoflurane and placed in a stereotaxic device. Anchor screws were placed above the cerebellum, right parietal hemisphere, and anterior cranium. The recording bundles designed to target prelimbic cortex, infralimbic cortex, medial amygdala, ventral hippocampus, primary visual cortex, mediodorsal thalamus, lateral habenula, lateral septum nucleus, nucleus accumbens, ventrolateral portion of the ventromedial hypothalamus, and orbital frontal cortex were centered based on stereotaxic coordinates measured from bregma. [Orbital frontal cortex: anterior/posterior (AP) 2.35mm, medial/lateral (ML) 1.0mm, dorsal/ventral (DV) from dura -2.75mm; infralimbic cortex and prelimbic cortex: AP 1.8mm, ML 0mm, DV -2.7mm from dura; medial amygdala: AP -1.25, ML 2.7mm, DV -4.3 from dura; lateral septum and nucleus accumbens: AP 1.0mm, ML 0mm, DV -4.0mm from dura; ventromedial hypothalamus, lateral habenula, and medial dorsal thalamus: AP -1.47mm, ML 0mm, DV -5.4mm from dura; ventral hippocampus and primary visual cortex: AP -3.0mm, ML 2.6mm, DV -3.0mm from dura]. We targeted infralimbic cortex and prelimbic cortex by building a 0.6mm DV stagger into the bundle. We targeted lateral septum and nucleus accumbens by building a 0.3mm ML and 1.5mm DV stagger into the bundle. We targeted lateral habenula, medial dorsal thalamus, and ventral medial hypothalamus by building a 0.3mm ML, and 1.9mm and 2.5mm DV stagger into our electrode bundle microwires. We targeted primary visual cortex and ventral hippocampus using a 0.3mm ML and 2.5mm DV stagger in our electrode bundle microwires. For open- and closed-loop optogenetic stimulation experiments, the addition of a Mono Fiberoptic Cannula coupled to a 2.5mm metal ferrule (NA: 0.22, 100µm [inner diameter], 125µm buffer [outer diameter], MFC_100/125-0.22, Doric Lenses, Quebec) was built into the medial prefrontal cortex bundle. The tip of the fiber was secured 300µm above the tip of the IL microwires. Mice were allowed 10-15 days for recovery from surgery before behavioral testing. For the medial prefrontal cortex projection-targeted physiological experiment, mice were implanted in medial prefrontal cortex (mPFC), and one of the three projection-targeted regions (nucleus accumbens [NAc], medial amygdala [MeA], or lateral habenula [LHb]) using the coordinates described above. A 2.5mm metal ferrule (NA: 0.22, 100µm [inner diameter], 125µm buffer [outer diameter], MFC_100/125-0.22, Doric Lenses, Quebec) was built into the microwire bundles targeting medial prefrontal cortex and the bundle targeting medial prefrontal cortex terminals in the projecting region.

### Viral surgery

For optogenetic experiments targeting mPFC soma, we used CD1 mice that showed an attack latency < 60s when exposed to an intact C57BL/6J (C57) male. 40 CD1 mice were anesthetized with 1% isoflurane and placed in a stereotaxic device. The mice were unilaterally injected with AAV2-CamKII-ChR2-EYFP (purchased from the Duke Viral Vector Core, Durham, NC; courtesy of K. Deisseroth), based on stereotaxic coordinates from bregma (left infralimbic cortex: AP 1.8mm, ML 0.3mm, DV -2.0mm from the brain). A total of 0.5µL of virus was infused at the injection site at a rate of 0.1µL/min over 5 minutes, and the needle was left in place for 10 minutes after injection. For the open-loop stimulation experiment, CD1 mice were implanted with an optic fiber (Mono Fiberoptic Cannula coupled to a 2.5mm metal ferrule (NA: 0.22, 100µm [inner diameter], 125µm buffer [outer diameter], MFC_100/125-0.22, Doric Lenses, Quebec)) 0.3mm above the injection site immediately after viral syringe was removed. These mice were allowed 3 weeks for recovery prior to behavioral testing. For the closed-loop experiments, 11 CD1 mice were allowed 3 weeks for viral expression prior to implantation with an optrode. For the nonsynchronous stimulation experiment, 19 CD1 mice were allowed 3 weeks for viral expression prior to implantation with an optic fiber with an electrode connector attached.

For the ESR1-Cre validation experiment, 13 ESR1-Cre+/CD1 F1 offspring were bilaterally injected with AAV2-hSyn-DIO-GqDREADD (obtained from Addgene) based on stereotaxic coordinates measured from bregma (AP -1.5mm, ML ±0.7mm, DV -5.7mm from the dura). A total of 0.3µL of virus was infused bilaterally at a rate of 0.1µL/min, and the needle was left in place for 5 minutes after injection. Two weeks after viral infusion, F1 males were screened for aggressive behavior towards females. The F1 males received i.p. injections of CNO (1mg/kg) at the start of the screening session. 35 minutes after injection, a novel C57 female was placed in the home cage for 5 minutes. Screening was repeated one week and two weeks later. Only F1 males who attacked females for at least two of the three screening sessions (9/13 mice) were implanted with electrodes ^18^. The eight mice that showed good surgical recovery were subjected to further experiments.

For mPFC projection-targeted behavioral experiments, we used 46 male CD1 mice that showed an attack latency <60s and initiated attacks at least three times within three minutes when exposed to a C57 male mouse (categorized as aggressive mice). These mice were unilaterally injected with AAV2-EF1a-DIO-ChR2-eYFP (obtained from Addgene) in the left mPFC based on stereotaxic coordinates measured from bregma (AP 1.8mm, ML 0.3mm, DV -2.0mm from the dura), and AAVrg-EF1a-Cre-mCherry (obtained from Addgene) was injected in the downstream target region based on stereotaxic coordinates measured from bregma (NAc: AP 1.0mm, ML 0.9mm, DV -3.8mm from the dura; MeA: AP -1.25mm, ML 2.75mm, DV -4.3mm from the dura; or LHb: AP -1.6mm, ML 0.4mm, DV -2.2mm from the dura). A total of 0.3µL of virus was infused in the mPFC and 0.3µL of virus was infused in the downstream target region at a rate of 0.1µL/min. The needle was left in place for five minutes after injection. Immediately after the viral syringe was removed from the downstream target region, mice were implanted with an optic fiber (Mono Fiberoptic Cannula coupled to a 2.5mm metal ferrule (NA: 0.22, 100µm [inner diameter], 125µm buffer [outer diameter], MFC_100/125-0.22, Doric Lenses, Quebec)) 0.3mm above the downstream target region injection site. Four weeks after viral infusion/optic fiber implantation, CD1 mice were tested for the behavioral effects of projection-targeted stimulation.

For mPFC projection-targeted electrophysiological experiments, six male CD1 mice that did not show aggression during screening were anesthetized with 1% isoflurane and placed in a stereotaxic apparatus. Anchor screws were placed over the anterior cranium, parietal cortex, and cerebellum. These mice were injected with AAV-CAG-DIO-ChR2-eYFP and AAVrg-EF1a-IRES-Cre-mCherry, as described above, to target mPFC → NAc (N=2 mice), mPFC → MeA (N=2 mice), or mPFC → LHb (N=2 mice). Optrodes targeting mPFC and the target regions were then implanted as described.

For the LinCx circuit-editing experiment, 60 retired breeder male CD1 mice were screened for aggressive behavior over three days for 5 minutes/day. Only mice that 1) attacked on days 2 and 3, 2) had an average total attack time >20 seconds, and 3) had an average latency to first attack <60 seconds were included in the experiment. Anticlust in R (R Foundation for Statistical Computing, Vienna, Austria, Ver. 4.4) was used to separate these mice into two groups (LinCx-edited mice and control mice) with equivalent attack and non-attack engagement times. Group assignments were performed independently across two cohorts of mice (N=7 and 5 control mice, and N=7 and 6 LinCx-edited mice for cohort 1 and 2, respectively). Mice in the LinCx-edited experimental group were then injected with 0.3µL AAV9-CaMKII-Cx34.7M1-T2A-mEmerald in left mPFC (AP 1.8 mm, ML 0.3 mm, DV -2.0 mm from dura) and 0.3µL AAV9-CaMKII-Cx35_M1_-T2A-mCherry in left NAc (AP 1.0 mm, ML 0.9 mm, DV -3.8 mm). Virus was injected at 0.1µL/min using a microinjection syringe pump (WPI Micro2T SMARTouch with UMP3 pumps). Syringe remained in place for at least 5 minutes after each injection. Control mice were injected with AAV9-CaMKII-EGFP (Addgene viral prep # 50469-AAV9, a gift from Bryan Roth) in mPFC and AAV9-CaMKII-mCherry (Addgene viral prep # 114469-AAV9, a gift from Karl Deisseroth) in NAc using the same procedure noted for the LinCx-edited mice.

### Histological analysis

Histological analysis of implantation sites was performed at the conclusion of experiments to confirm recording sites and viral expression (Supplementary Figs. S11-S12, and S14). Animals were perfused with 4% paraformaldehyde (PFA), and brains were harvested and stored for at least 96 hrs in PFA. Brains were cryoprotected with sucrose and frozen in OCT compound and stored at -80°C. Brains were later sliced at 40µm. Brains from mice used to train and validate the network were stained using NeuroTrace fluorescent Nissl Stain (N21480, ThermoFisher Scientific, Waltham, MA) using standard protocol. Specifically, Nissl staining for brain tissue occurred on a shaker table at room temperature. Tissue was washed in PBST (0.1% Triton in phosphate-buffered saline solution) for 10 minutes. It was then washed for 5 minutes in PBS twice. The tissue was then protected from light for the remainder of the protocol. The tissue was incubated in 1:300 Nissl diluted in 2 mL PBS for 10 minutes. After the Nissl incubation, tissue was washed once in 0.1% PBST for 10 minutes, then twice in PBS for 5 minutes. Brains from ESR1 mice and mice used for 20 Hz open- or closed-loop stimulation were mounted in Vectashield mounting medium containing DAPI (H-1200-10, Vector Laboratories, Newark, CA). Images were obtained at 10x using an Olympus fluorescent microscope.

### Neurophysiological and video data acquisition

Mice were connected to a Cerebus data acquisition system (Blackrock Microsystems, UT, USA) while anesthetized with 1% isoflurane. Mice were allowed 60 minutes in their home cage prior to behavioral and electrophysiological recordings. Local field potentials (LFPs) were bandpass filtered at 0.5-250Hz and stored at 1000Hz. An online noise cancellation algorithm was applied to reduce 60Hz artifact (Blackrock Microsystems, UT, USA). Neural spiking data was referenced online against a channel recording from the same brain area that did not exhibit a SNR>3:1. After recording, cells were sorted using an offline sorting algorithm (Plexon Inc., TX) to confirm the quality of the recorded cells. Only cell clusters well-isolated compared to background noise, defined as a Mahalanobis distance greater than 3 compared to the origin, were used for the unit-electome network correlation analysis. We used both single (well-isolated clusters with ISI<1.5) and multi-units (well-isolated clusters with ISI>1.5; N=186 total neurons) for our analysis as our objective was to determine whether electome network activity was reflective of cellular activity. Neurophysiological recordings were referenced to a ground wire connected to anchor screws above the cerebellum and anterior cranium. Video data was acquired using Neuromotiv (Blackrock Microsystems, UT, USA) and synchronized with neural recordings.

### Behavioral recordings and analysis for training/testing models

The 45 CD1 mice used for training and testing the electome network model were first subjected to screening to assess their basal level of aggressiveness. Screening occurred once a day for three consecutive days prior to surgical implantation. Animals were screened in cohorts. For each screening session, an intact male C57 was placed in the CD1’s home cage for 5 minutes and the latency to first attack was recorded. To ensure that our network generalized broadly across CD1 mice, we used a training and testing set for which ∼50% of the mice showed high aggression during screening (i.e., latency to attack < 60s), and ∼50% of the mice showed low to moderate aggression (i.e., latency to attack > 60s). Animals that showed no aggression at all during screening (16/45 mice) were excluded from further experiments.

All screening/testing occurred within the home cage of mice except for the quantification of cortical stimulation-induced gross locomotor activity. These latter experiments were performed in a 44cm × 44cm × 35cm (L×W×H) open field arena. Subject mice (CD1 and ESR1 males) were acclimated to the recording tether for three days prior to the first recording session. Each acclimation session involved anesthetizing the mouse with 1% isoflurane, tethering the subject mouse, allowing 60 minutes to recover from isoflurane, then placing a male C57 in the home cage for 5 minutes. Mice were then anesthetized with isoflurane again and detached from the tether. The aggression level of experimental mice was determined based on average latency to attack intruder mice during the second and third acclimation sessions.

After screening, 29 mice were implanted, and data was acquired across 1-6 behavioral testing/recording sessions following recovery. Sessions were separated by 5-7 days. Recordings for all social encounters were performed in the home cages of the CD1 mice. Each behavioral testing session began with 5 minutes of recording prior to introduction of the social stimulus. All mice were subjected to encounters with an intact C57 male mouse and a female C57 mouse. A subset of 18 CD1 mice were also subjected to an encounter with a castrated male mouse. For this selection, the aggressiveness of each mouse was defined as the median latency to an attack an intact male intruder across the six initial recording sessions. The eight most aggressive mice were also subjected to exposure to objects covered in CD1 mouse urine. Object pairs (one clean and the other covered in urine) included yellow duplex blocks, curved red duplex blocks, weighted 5 mL conicals, glassware tops, and objects assembled from black Legos®. The CD1 mice were exposed to a different pair of objects during each session. Order of exposure to stimulus mice and objects was shuffled for every session. Six CD1 mice were recorded under all four conditions. Data observations (1 second each) were pooled across 20 CD1 mice for training the network model.

For ESR1-Cre+/CD1 male behavioral testing, eight mice were injected with either saline or CNO (1mg/kg, i.p.) after the five-minute baseline recording. 35 minutes after this injection, mice were exposed to an intact male C57, a castrated male C57, and a female C57, presented in pseudorandom order. Mice were subjected to six total recording sessions, three in which they were treated with saline and three in which they were treated with CNO, again in pseudorandom order. Sessions were separated by 5 days to allow an adequate washout of CNO^53^.

Behavior was scored for each second as an "aggressive behavior", "non-aggressive social interaction", or "non-interaction". One-second windows were identified as "aggressive behavior" if the mouse was engaged in biting, boxing (kicking/clawing), or tussling behavior ^34^. Windows were labeled as "non-aggressive social interaction" if the mouse had his nose or forepaws touching the stimulus mouse (intact male/female/castrated male) or object, but was not biting, boxing, or tussling. Examples of behaviors labeled as "non-aggressive social interaction" included sniffing, grooming, or resting (placing nose or forepaws against the subject mouse, but not moving). If the stimulus mouse had his/her forepaws or nose on the CD1 but it was not reciprocated, this was labeled as "non-interaction". In our experimental settings, CD1 straight approach, sideways approach, and chasing of the stimulus mouse could result in aggressive behavior (biting/kicking/tousling), non-aggressive social interaction (nose or paw touch), or withdrawal without any touch. Thus, while sideways approach and chasing are regularly labeled as "aggressive" in the literature, and straight approach is not^34, 54, 55^, these behaviors lacked consistent resolutions. Moreover, mice also demonstrated these behaviors towards female and castrated mice (i.e., in non-aggressive social behavior contexts). Therefore, one-second windows containing these behaviors were labeled as "non-interaction". All other timepoints not labeled as "aggressive behavior" or "non-aggressive social interaction" were also labeled as "non-interaction". These behavioral criteria were selected to include ethologically aggression-related behaviors and maximize the likelihood that the CD1 was aware of the presence of the stimulus mouse or object during the behavioral window, while remaining confident in the classification of "aggressive behavior" and "non-aggressive social interaction" window labels.

Tail rattling is well-recognized in the literature as an aggressive behavior^34^, and it was consistently only demonstrated by aggressive mice towards intact male mice in our experimental settings. Thus, we included this behavior in the "aggressive behavior" category. In our subset of 20 mice used for training the network, tail rattling was observed 8 ± 4s out of the 135 ± 26s "aggressive behavior" windows per mouse.

The videos used to generate the behavioral labels for training and testing our machine learning model were hand-scored by a trained researcher. Videos from ESR1-Cre+/CD1 mice and optogenetic stimulation were automatically tracked using DeepLabCut ^56, 57^. This information was then used for creating behavioral classifiers in SimBA ^58^.

### LFP preprocessing and signal artifact removal

Each LFP signal was segmented into 1s non-overlapping windows. If there were multiple intact channels implanted in a region, they were averaged to produce a single signal. Windows with non-physiological noise were removed using an established automated heuristic ^8^. Briefly, the envelope of the signal in each channel was estimated using the magnitude of the Hilbert transform. The Median Absolute Deviation (MAD) of the magnitude was then calculated on each channel of each recording. Signal was marked as non-physiological if the envelope exceeded a high threshold (5x MAD, which is roughly 4x the standard deviation for a normally distributed signal). Any data adjacent to non-physiological data that had an envelope value above a smaller threshold (0.167 MAD) was also considered non-physiological. All data marked in this way was ignored when averaging channels for each region. Any channels with a standard deviation less than 0.01 were removed as well. If no channels were usable for a given region within a window, that whole window was removed from the data. This set of heuristics resulted in 34.7±5.1% of the data windows being excluded from analysis. After this, 60Hz line artifact was further removed using a series of Butterworth bandpass filters at 60Hz and harmonics up to 240Hz with a stopband width of 5Hz and stopband attenuation of -60dB. Finally, the signal was downsampled to 500Hz.

### Estimation of LFP oscillatory power, coherence, and directionality

Signal processing was performed using MATLAB (The MathWorks, Inc. Natick MA, Version 2023a). For LFP power, spectral power was estimated using Welch’s method using a 1-second window and 1-second steps at a 1Hz resolution. Coherence was estimated pairwise between all recorded regions using Welch’s method and magnitude-squared coherence defined as

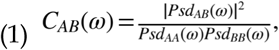

where and are two regions and *Psd_AA_(ω)* and *Psd_AB_(ω)* are the power and cross-spectra at a given frequency , respectively.

Directionality features -- a correlate of directional transfer of information across a brain circuit -- were estimated using the Multivariate Granger causality (MVGC) MATLAB toolbox ^59^. To get stable Granger causality estimates, a 6^th^-order highpass Butterworth filter – with a stopband at 1Hz and a passband starting at 4 Hz – was applied to the data using the *filtfilt* function (MATLAB, The MathWorks, Inc. Natick MA). Granger causality values for each window were estimated with a 20-order AR model at 1 Hz intervals to align with the power and coherence features. These directionality features were processed identically to a previously reported approach ^8^. Briefly, directionality features were exponentiated to approximately maintain the additivity assumption made implicitly by Nonnegative Matrix Factorization (NMF) models ^8, 60^ as *exp(f_A→B_(ω))* where *f_A→B_(ω)* is the Granger causality at frequency from region *A* to region *B*. The exponentiated feature is a ratio of total power to unexplained power. Exponentiated directionality feature values were truncated at 10 to prevent implausible values.

### Data for single-region and network-level machine learning analyses

We used 21460 seconds of behavioral/neural data, pooled across the 20 mice to train/validate our single-region and network models. This included a total of 4680 seconds while mice were socially isolated in their home cage, 14890 seconds when CD1 mice exhibited non-aggressive social behavior (3542 seconds towards intact males, 9067 seconds towards females, and 2281 seconds towards castrated), and 1890 seconds when mice exhibited aggressive behavior towards the intact males.

### Discriminative Cross-Spectral Factor Analysis – Nonnegative Matrix Factorization

Our machine learning approach was designed to distinguish between behavioral windows when the CD1 mice showed aggressive behavior towards intact C57 males and windows when they exhibited pro-social behavior (interactions towards intact C57 males, castrated C57 males, or C57 females), and it was designed also to model neural variance. Here, we used neural and behavioral data from 29 mice to establish the final model, with a split of 20 and 9 mice for model training and testing, respectively.

Our approach used the Discriminative Cross-Spectral Factor Analysis – Nonnegative Matrix Factorization (dCSFA-NMF) model ^38^. This approach assumes that each window is an independent stationary observation and examines dynamics in brain activity only at the scale of windows. In this case, the relevant dynamics happen at the scale of 1-second windows. A 1-second window was chosen to balance capturing fine-grained transient behavior with sufficient length to properly estimate spectral features ^8^. Each window has associated spectral power, coherence, and directionality features (in total features), which is represented as for the window. Each window (Fig. 1e, top; Supplementary Fig. S3a, top) was associated with a behavioral label that identified which condition the CD1 mouse was subjected to during that window (intact male, castrated male, or female) and whether the CD1 mouse was engaged in aggressive or non-aggressive behavior during that window (coded as y_i_ ∈ (0, 1})

As a short description of the dCSFA-NMF model, all spectral power, coherence, and directionality features are described as an additive positive sum of k nonnegative electrical functional connectome (electome) networks. This model is learned using a supervised autoencoder. The objective we use to learn the parameters is

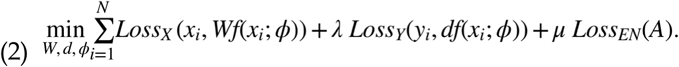

Here, Lossx is the reconstruction loss of the features derived from electrophysiology, which for our work was an L_2_ loss. Our predictive loss Lossy is the cross-entropy loss commonly used for binary classification. Each of the k networks is represented in vector form and combined to make a matrix W Rpk, where each column will represent an electome network. This matrix can also be thought of as a decoder network that is limited to a single linear layer in an autoencoder framework. The electome network activities are given by the multi-output encoder function f(x;φ):R^p^R^k^, where φ represent the parameters of the function. These activities correspond to how strongly expressed a network is within a window, allowing for variations in network strength over time. In our model, the multi-output function was an affine transformation of Ax+B followed by a softplus rectification, defined as softplus(x)= log(1+expx), thus = {A,b} . Parameters dR^k^ represent the relationship between the electome networks’ activities and the behavior. A sparsity constraint is enforced so that d=[d_1_,0, 0], meaning that only a single electome is used to predict aggressive or non-aggressive behavior, simplifying interpretation (Fig. 1e, bottom left; Supplementary Fig. S3a, bottom). The first electome network is referred to as the supervised network, since it is used to predict the aggressive state, and the rest are referred to as unsupervised electome networks, as they are not directly linked to the behavior of interest. λ is a weighting parameter used to control the relative importance of prediction and reconstruction. In other words, this term varies how much the training values predicting aggressive or non-aggressive behavior compared to explaining the LFP feature variance. We chose a value of λ =1 , which kept the two losses approximately equal during training.

Previous work has also found that the reconstruction loss can reduce overfitting and make the learned predictions more robust ^61^. To further reduce overfitting of the predictive aspect of the network activities, we applied an elastic net loss^62^, on the encoder parameter A that produce the network activities. Loss_EN_ has a weighting μ and the ratio between the L1 and L2 losses set to .5. The value for μ was set to a small value that had worked well previously ^63^. In this work, power features were also upweighted by a factor of 10 to accommodate that there were many more coherence and directionality features and truncated at 6 to prevent outliers from dominating the predictions.

These models and statistical analyses were implemented with Python 3.7 and Tensorflow version 2.4. Parameters were learned with stochastic gradient descent using the Nesterov accelerated ADAM optimizer ^64^. Learning was performed for 30000 iterations, which was observed to be ample for parameter convergence. The learning rate and batch size were set to 1e-3 and 100 respectively, values that have empirically performed well in similar applications ^63^.

Predictive performance of the electome network was evaluated using data from new mice that were not included in learning the electome network. Network scores were estimated as an evaluation of the encoder learned during training of the dCSFA-NMF model.

### Hyper-parameter selection

The dCSFA-NMF procedure requires selection of several settings in the algorithm. Specifically, we must choose: the number of electome networks ; the importance of the supervised task ; the relative importance of the power features, coherence features, and directionality features; and the parameterization of the mapping function , which is referred to as the encoder. Besides , these settings were chosen to match previously used values or to follow heuristics. Specifically, in our prior work, we demonstrated that the inferred model is highly insensitive to λ ^63^. Thus, we chose a λ value to give roughly equal weight to the predictive and generative tasks. Similarly, since the number of power features grows linearly and the number of coherence and directionality features grows quadratically with the number of brain regions (as they are pairwise), we chose to weight the power features to roughly match the coherence features. Since the decoder is chosen to be linear (given by the matrix W above), as noted above we also chose a linear mapping function for the encoder followed by a softplus function to ensure non-negativity. This approach served to limit complexity.

To choose the value of K, we evaluated the reconstruction error (Mean Squared Error) on the 20 training mice, which determined how well the electome networks described the neural features, how consistent the electome networks were, and how well they predicted the behavioral state (aggressive vs. non-aggressive) as we varied the total number of electome networks K while using a single supervised network (e.g., 1 supervised network and K-1 unsupervised networks). We did not expect the prediction quality to be especially dependent on the number of electome networks, as our previous work has demonstrated that K is not an especially important parameter in terms of predictive performance ^63^, but that it can impact reconstruction error and total model fit.

For our main selection criteria, we used the Bayesian Information Criterion (BIC) to select the number of unsupervised networks to use in the final network model. The BIC is defined as:

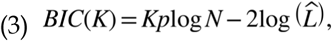

where kp is the number of model parameters (p is the number of spectral features), N is the number of samples, and L_is_ is the likelihood of the observed data using all estimated model parameters (φ,w,d) . This criterion balances the model fit quantified bylog(L) with the complexity quantified by kplog_N_ . In this work, -log(L) is an L_2_ loss, corresponding to a Gaussian observational likelihood, as used in the training above. Using the training animals above, the model parameter were learned from 80% of the data while model parameter L was evaluated on a 20% hold-out set to avoid overfitting. The BIC was evaluated for all dimensionalities from 1-20 networks, and the lowest value was selected as the best model. Since 7 unsupervised networks provided the best fit (a BIC of 5457701, see also Supplementary Fig. S2b), our final full network model utilized a total of 8 networks, 1 supervised and 7 unsupervised, across all 11 regions.

For each single-region model, we trained 3 unsupervised networks and a single supervised network. Here, we reduced the number of networks compared to the full network model, given the dramatic reduction in the number of covariates considered by the model, as there are no longer pairwise features calculated between regions. Our objective was to compare the predictive performance of the single-region models against each other and the full network model. Since the predictive performance is driven by the supervised network^63^, the smaller latent dimensionality of the single-region models had no impact on our final conclusions.

### Single-region decoding of social behavioral state

To test the efficacy of any single brain region as a biomarker for aggression, we extracted power at 1-Hz frequency bins over 1-56 Hz from each region. One-second windows were pooled from the 20 CD1 training mice to generate a series of dCSFA-NMF models for each of the 11 brain regions (see above section for model hyperparameters). The models were trained to distinguish behavioral windows of one among three social states exhibited by CD1 mice: 1) male-directed aggressive behavior, 2) female non-aggressive social interactions, and 3) castrated male non-aggressive social interactions. We also developed a model to distinguish 4) periods when CD1 mice were isolated in their home cage from any of the three social states. Each model was then tested on data from a hold-out set of nine mice. The area under the receiver operating curve (AUC) was calculated for each hold-out mouse to determine the performance of the model. False discovery rate was used to correct for multiple hypothesis testing.

### Sensitivity of supervised electome network parameters to total number of electome networks

A frequent concern of latent variable models, including dCSFA-NMF, is the dependence of the network parameters *W* and encoder parameters *A* on the choice of latent dimensionality *K*, which determines the total number of electome networks in our models. In particular, in our case, we are most interested in how robustly we learn the encoder parameters and network parameters related to the single supervised network. To address this concern, we performed a sensitivity analysis to determine the extent to which the choice of *K* influenced the supervised electome network parameters (first column of *W*) and encoder (first row of A). In this sensitivity analysis, we estimated a dCSFA-NMF model allowing the number of total networks to range from 2-20 while keeping a single supervised network. We then compared the similarity between each learned supervised encoder and supervised electome network parameters to our model with eight networks (the final, full model used in this work that has 1 supervised and 7 unsupervised networks). This was quantified using the cosine similarity, which measures the angle between two vectors, ranging from -1 to 1. A value of 1 indicates perfect alignment (pointing in the same direction), 0 is orthogonal, and -1 indicates that the vectors point in opposite directions) (Supplementary Fig. S2a). We applied this to the vectors that define the supervised electome network (also referred to as a decoder due to the autoencoder structure; first column of *W*) and the supervised encoder (first row of *A*).

The supervised encoders were virtually identical across all the models except the one that utilized three networks, which learned a network that was positively associated with aggression (Supplementary Fig. S2a). We found that the supervised network parameters (decoder) maintained a strong consistency across most dimensions, particularly between 5-10 networks, as shown by the cosine similarities being greater than 0.95.

To evaluate how large these cosine similarities are, we created a null distribution of the similarities across randomly chosen generative networks, where we randomly matched networks, meaning that we randomly matched the rows of *W* corresponding to network parameters (decoder) and columns of *A* corresponding to a single randomly chosen network (encoder). These similarities were substantially lower. In summary, across a range of tested latent dimensionalities, the supervised electome model converged on highly similar behavior-linked network patterns. In other words, varying the number of latent components did not materially change the structure, regional composition, or behavioral associations of the learned electome network. This suggests that the aggression-related network is a stable, data-driven feature of the recordings rather than an artifact of selecting a particular model dimensionality. Importantly, this conclusion applies to the supervised electome framework (that is, networks constrained by behavioral prediction), rather than to purely unsupervised decompositions.

### Single-cell correlation to electome network activity

Data acquired during the third behavioral testing session was from the 20 implanted training mice were used for cellular analysis. We used Spearman correlation to quantify the relationship between cellular firing windows and the activity of the electome network used to classify aggressive behavior. We performed 1000 permutations by randomly shuffling 1-second windows within each class for aggressive behavior and non-aggressive social interactions with male, female and castrated C57 mice. This approach maintained the relationship between network activity and behavior and the relationship between cell firing and behavior. We then calculated the Spearman correlation between network activity and cell firing for each permutation. A cell was deemed positively correlated if its unshuffled Spearman Rho was above 97.5% of the permuted distribution and negatively correlated if it was below 2.5%.

### High-density single unit recordings

Three mice which showed aggression during screening were anesthetized with isoflurane (1%) and placed in a stereotaxic device, and metal ground screws were secured to the anterior cranium and above cerebellum. Multi-wire electrodes were implanted to target OFC, MeA, VHip, V1, VMHvl, MDThal, and LHb based on the coordinates noted above, and a 1024-channel silicon probe (NeuroNexus, SINAPS_4S_1024) was implanted to target IL, PL, NAc, and LSN (-1.7mm AP, centered at the midline, measured from bregma; - 4mm DV from the dura). Individual shanks were spaced by 500µm on the probe such that the front two targeted medial prefrontal cortex bilaterally at ±0.25mm ML. The multi-wire electrode implant and silicon probe were secured to the same ground.

Following surgical recovery, mice were habituated to the recording cables for 1-2 weeks and tested again for aggression. One of the implanted mice failed to show aggression and was removed from further testing. Neural data was recorded from the other two implanted mice during exposures to intact male, castrated male, and female mice using the Cerebus System (Blackrock Microsystems, UT, USA), as described above. Neural data was sampled concurrently from the implanted silicon probe at 20kHz using a SmartBox Pro acquisition system (NeuroNexus). An analog timing signal, consisting of 10ms pulses with a pseudorandomized inter-pulse-interval of 8-24 seconds, was generated using the analog output from the Cerebus system, and it was stored at 1000Hz alongside recorded neural data via the analog input channel on the SmartBox Pro. This timing signal, recorded concurrently by both systems, was used to align the two sets of neural data for analysis.

To extract single unit activity, neural activity was converted to the *Neurodata Without Borders (NWB)* format, high-pass filtered at 300Hz, and the medial 1024 channels were automatically sorted with Kilosort4^65^. Single units were identified based on ISI violations < 0.5, presence ratio > 0.9, an amplitude cutoff < 0.1 (473/1052 total sorted cells)^66, 67^.

### Real-time encoder approximation for closed-loop optogenetic experiments

Because calculating all the directionality features was too computationally demanding for real-time calculation, we developed a faster dCSFA-NMF model that relied only on power and coherence features for estimation of *EN-AggINH* to use in the closed-loop stimulation experiments. This ’fast’ model was trained on the same data used to learn the full network model. The model was trained using regularized regression to best predict the output of the full encoder,f(x_i_: φ), defined above. As such, this reduced encoder is also an affine transformation followed by a softplus activation with a smaller parameter set, φ_r_= {A_r_,b_r_}. The only difference from the full encoder is that the reduced encoder uses a smaller number of neural features and is otherwise the same mathematical form. This approximation explained a large component of the variance of the supervised network score derived from the hold-out validation mice (R, p-value <10^-16^).

### Optogenetic stimulation

Mice were anesthetized with 1% isoflurane, then tethered to an optic patch cable placed over the optic fiber cannula. The mice were then allowed 60 minutes for recovery prior to behavioral testing and undergoing two stimulation sessions. For each session, mice were exposed to intact male and female mice. A subset of mice was also exposed to a castrated male mouse. During each condition, the CD1 mouse received two segments of alternating yellow (593.5nm, OEM Laser Systems, Model No. MGL-F-593.5/80mW) and blue (473nm, Crystal Laser LC, Reno, NV. Model No. DL473-025-O) light stimulation, for two minutes each. During the first session, mice were stimulated with yellow light first (yellow, blue, yellow, blue), and during the second session mice were stimulated with blue light first (blue, yellow, blue, yellow). CD1 mice received light stimulation for the entirety of the two-minute segment. CD1 mice received light stimulation for the entirety of the two-minute segment.

For closed-loop optogenetic stimulation, mice anesthetized with 1% isoflurane were tethered to an optic patch cable and connected to the recording system. The mice were then allowed 60 minutes for recovery prior to behavioral testing/neural recordings. Mice experienced three sessions of behavioral testing followed by two sessions of closed-loop stimulation. Stimulation sessions were separated by 5-7 days. For each of the behavioral testing sessions, CD1 mice were exposed to intact C57 males, females, and castrated male mice for 5 minutes each. Testing sessions two and three were used to determine a reduced network threshold at which 40% of aggressive behavioral windows could be detected. For each stimulation session, mice were recorded for 3 minutes of baseline in their home cage, then during the three social encounters. The order of the three social encounters was shuffled for each session. Mice were recorded in an open field for 5 minutes after each session. During each social condition, the CD1 mouse went through two-minute segments of potential blue light stimulation (473nm, Crystal Laser LC, Reno, NV. Model No. DL473-025-O), then yellow light stimulation (593.5nm, OEM Laser Systems, Model No. MGL-F-593.5/80mW), and blue and yellow light stimulation again. During these two-minute segments, mice only received stimulation for one second when the reduced network score dropped below threshold. Behavior was averaged across the two stimulation sessions.

For nonsynchronous stimulation, each CD1 mouse was pseudorandomly matched to a different mouse that had been used for closed-loop stimulation. Each nonsynchronous mouse was then subjected to the identical order of social conditions and blue and yellow light stimulation blocks as their individually matched closed-loop mouse. Light stimulation was delivered using the pattern implemented for the matched mouse receiving closed-loop stimulation.

For the projection-targeted-stimulation behavioral experiment, CD1 males were exposed to four-minute segments of light stimulation in the following sequence: yellow, blue, yellow. Mice received light stimulation for the entirety of each segment. Within each stimulation segment, an intact male C57 and a female C57 were placed in the CD1 cage for two minutes each in pseudorandom order. A subset of the mice were also exposed to a castrated male.

Immediately prior to experiments, light levels were calibrated using a power meter (ThorLabs, Model No. PM100D) and delivered using a Waveform generator (Agilent Technologies, Model No. 33210A) for the open-loop experiment and projection-targeted-stimulation behavioral experiment. For the closed-loop and nonsynchronous stimulation experiments, as well as the projection-targeted-stimulation physiological experiment, the laser was activated using analog output from the Cerebus recording system.

### Behavioral analysis for optogenetic experiments

For open-loop, closed-loop, and nonsynchronous stimulation optogenetic stimulation experiments, mice were exposed to intact male mice, castrated male mice, and female mice in pseudorandomized order. Mice were stimulated across two testing sessions and the time mice engaged in aggressive and non-aggressive behaviors was averaged across the two yellow light stimulation segments and compared to the time mice engaged in these behaviors during the two blue light stimulation segments. For the mPFC projection-targeted experiment, mice were exposed to intact male mice and female mice in pseudorandomized order. A subset of mice was also exposed to a castrated male. Mice were stimulated during a single session. The amount of time mice spent engaged in aggressive and non-aggressive behaviors were averaged across the two yellow light stimulation segments and compared to the time mice engaged in these behaviors during the blue light stimulation segment.

### Mediation Analysis

For the Baron and Kenny approach ^46^ to test whether *EN-AggINH* expression mediated the behavioral effect of neurostimulation, we first established the two prerequisite relationships: (i) that neurostimulation significantly altered *EN-AggINH* expression (Figure 5d) and (ii) that neurostimulation significantly reduced aggressive behavior (Figure 6a, right). We then tested whether inclusion of *EN-AggINH* expression attenuated the direct effect of stimulation on behavior, consistent with mediation. For the closed-loop optogenetic stimulation experiment, we focused on windows of LFP data when either the blue (treatment condition) or yellow laser (control condition) was activated to match the conditions between the treatment and control as closely as possible, which is necessary for clean statistical analysis. We followed the procedures outlined in the LFP preprocessing and signal artifact removal methods section to preprocess the data and remove data with significant artifacts. *EN-AggINH* expression was calculated by projecting the neural features into the learned model. The remaining data after the matching and preprocessing was then fit into two logistic regression models to predict aggressive vs. non-aggressive behavior using the statsmodel package in Python. ^68^. The first model only used a constant and whether the subject had received blue or yellow light stimulation in that time window to predict aggressive vs. non-aggressive behavior in a logistic regression for each window of data, and the second model added *EN-AggINH* network expression during that window as an additional covariate. These two models were compared by using a likelihood ratio test to evaluate whether the second model was significantly better at predicting aggressive vs. non-aggressive behavior.

For the causal mediation analysis, we used the same data as described above in the classic mediation analysis. We define the treatment as blue versus yellow light stimulation (control), the mediator as *EN-AggINH* expression, and the outcome as aggressive versus non-aggressive behavior. These data were then used in the causal mediation analysis approach proposed by Kosuke, Keele, and Tingley ^47^ by using the statsmodels package in Python ^68^.

### Physiological assessment of mPFC projection-targeted stimulation

Non-aggressive CD1 mice were implanted to target mPFC and one of three downstream regions as described above (NAc, MeA, or LHb). Mice were then connected to a fiberoptic patch cable in their home cage. Mice were subjected to two stimulation conditions: blue light delivered either to the mPFC soma or to the terminals of mPFC afferents within the downstream region for which each mouse was implanted. Mice were then stimulated with 40 light pulses (10ms, 1mW) at each location with a pseudorandom inter-pulse-interval between 8-24 seconds. Neural data was collected during stimulation as described above and subjected to a 4th order Butterworth bandpass filter from 0.5Hz to 60Hz. LFP data was then aligned relative to each light pulse and averaged across the 40 light pulses to yield a light-evoked LFP average (i.e., evoked potential) for mPFC and each downstream site. As a negative control, experiments were also performed with 40 pulses of yellow light (10ms, 1mW).

We quantified the evoked potential within the 1-60ms window and determined that light stimulation (blue or yellow) induced neural activity if the lowest value within this window was lower than activity within the 100-1000ms window prior to light stimulation and the 500-1000ms window following light stimulation. This corresponds to an alpha threshold of 0.05, Bonferroni-corrected for 60 comparisons. Blue light stimulation of mPFC soma and mPFC terminals induced neural activity for all three circuits tested. No significant evoked potentials were observed for any of the circuits in response to yellow light stimulation.

### Statistical comparison of directionality features within EN-AggINH

To compare the extent to which PL/IL → NAc, PL/IL → MeA circuits and PL/IL → LHb circuits were suppressed during aggression, we quantified the relative LFP spectral energy (RLE) as a measure of relative directionality for each circuit within *EN-AggINH*. This RLE is the weight of each directionality feature of *EN-AggINH* normalized to the sum of the weight of that feature across all the learned electome networks. To make a reasonable approximation to the confidence intervals on the relative weights of PL/IL → NAc, PL/IL → MeA, and PL/IL → LHb at 20 Hz, as shown in Figure 6c, we performed the following bootstrapping procedure: first, we fixed the non-relevant components of the inferred full electome model, and then found the best fit on the *EN-AggINH* parameters by bootstrapping over animals for 1000 repeats. While this uncertainty approach does not hold for machine learning models in general, we note that this approach is reasonable here given the mathematical structure of the machine learning model. In particular, the model loss is biconvex over the parameters with penalty terms, meaning that if the network activities are fixed the network parameters are strongly convex and the optimization will yield a single, unique optimal solution to *EN-AggINH* parameters. Hence, this approach allowed us to get reliable and reproducible estimates of uncertainty through bootstrapping. Under this approximate procedure, we found that in all repeats that the PL/IL → NAc relationship was stronger than the PL/IL → MeA, which was in turn stronger than PL/IL → LHb. This implies a p-value of <1e-4 under this approximate procedure. Thus, both the NAc- and MeA-targeted circuits are statistically stronger than the LHb-targeted circuit.

### Data exclusions

For the open-loop stimulation experiment, we behaviorally screened 30 CD1 mice. 10 mice that showed aggressive behavior towards intact males were implanted with stimulating fibers. Two mice were removed from behavioral analysis due to poor ChR2 expression.

To discover *EN-AggINH*, we behaviorally screened 45 CD1 mice and implanted 31 of these mice that showed aggression. Two of these mice were removed from further analysis because they failed to show aggression post implantation. Of the 297 total implantation sites in the remaining 29 training/validation and testing set of mice, 17 were mistargeted (∼5.7% error rate). Of these mistargeted implants, 13 were within 200µm of the targeted structure. Given our prior work demonstrating high LFP spectral coherence (in the 1-55Hz frequency range) across microwires separated by 250µm in both cortical and subcortical brain regions ^42^, we chose to retain these 13 implants in our analysis. The other four mistargeted implants were within 350µm of the targeted structure. The most reliably mistargeted site was ventral medial hypothalamus, for which 4 animals were implanted within 200µm of the target, and 2 animals were implanted within 350 µm of the target.

Machine learning analysis typically benefits from larger data sets. Thus, we concluded that maintaining a higher number of data points likely outweighed the effect of a small number of mistargeted brain regions, particularly since our LFP measures were robust to the targeting inaccuracies we observed histologically. As such, we pooled data from all 20 implanted animals to learn our initial model (the training cohort). We employed a similar strategy for our validation analysis, where an animal was only removed from the validation set if there was clear histological confirmation of mistargeting >200µm for any of the recorded regions. Specifically, presuming accurate targeting with 94.3% certainty and targeting within 200µm at a higher certainty, we included animals with missing or damaged histological slices in our analysis. However, if there was clear histological confirmation of mistargeting for any of the recorded regions (as was the case for 1 mouse), the animal was removed from the validation testing. Importantly, our validation procedure implies that the machine learning models were robust, regardless of any slight imprecision in the animals we utilized for training.

For the multi-wire and silicon probe recordings, we implanted three mice that showed aggressive behavior towards intact males. We removed one mouse because it showed poor aggression after implantation when tethered. For the ESR1-Cre+/CD1 experiment, we screened 13 mice across three sessions following treatment with CNO. The three testing sessions were performed over 3 weeks. Only the nine male mice that attacked females during two of the three sessions were implanted with electrodes ^18^. The eight mice that showed good surgical recovery were subjected to further experiments. All of these mice were retained for behavioral and physiological analysis.

For the closed-loop stimulation experiment, we screened 30 CD1 mice and implanted the eleven mice that showed aggressive behavior towards intact males. Two mice were removed from behavioral/physiological analysis due to poor ChR2 expression or electrode mistargeting. For the nonsynchronous stimulation experiment, we screened 40 CD1 mice and implanted the 19 mice that showed aggressive behavior towards intact males. Five animals were removed from final analysis due to poor tissue expression of ChR2.

For the mPFC-projection-targeted stimulation experiment, we screened 120 CD1 mice. The 46 animals that showed high aggression were injected with virus targeting mPFC → NAc (N=12), mPFC → MeA (N=16), or mPFC → LHb (N=18). 19 mice were removed from subsequent behavioral analysis because they showed low aggression during the first yellow light stimulation segment (i.e., less than 20s of aggressive behavior towards intact males), poor expression of ChR2, or mistargeting (mPFC → NAc: N=3/12; mPFC → MeA: N=7/16; mPFC → LHb: N=9/18). We used six mice that showed low aggression for the projection-targeted physiological experiments. Of these six mice, one mouse that was infected (virus) and implanted (electrodes) to target the mPFC → LHb circuit died prior to stimulation. One mouse that was infected (virus) and implanted (electrodes) to target the mPFC → MeA circuit was excluded because of poor ChR2 expression. To ensure balance observations across the three groups, we pseudorandomly removed one mouse from the mPFC → NAc circuit group.

For the LinCx-editing experiment, we screened 60 CD1 mice. 25 mice that showed aggressive behavior towards intact males were pseudorandomly assigned to the LinCx-edited (N=13) or control groups (N=12) to balance aggressive behavior and non-aggressive social behaviors between the groups. Mice were behaviorally screened a second time to ensure the robustness of these group assignments. Mice were then injected with virus and subjected to behavioral testing at 2, 5, and 8 weeks after surgery. One control mouse and one LinCx-edited mouse were removed from the final behavioral analysis because they showed low aggression (i.e., less than 20s of aggressive behavior towards intact males) during the second presurgical screening. Two additional LinCx-edited mice were removed from behavioral analysis: one mouse showed poor Cx34.7M1 expression in the mPFC, and the other showed bilateral spread of Cx35_M1_ to mPFC. We retained one control mouse that showed poor expression of mCherry in NAc in our analysis since we did not anticipate that mCherry expression in the NAc would alter social behavior. Importantly, presurgical aggressive and non-aggressive social behavior remained balanced between the LinCx-edited and control groups with these four mice removed (see Fig. 6h, N=10 LinCx-edited mice and N=11 control mice).

### Statistics

GraphPad Prism 10 and MATLAB 2023a were used for statistical analyses of behavior and network activity. Parametric testing was used for comparison of behavioral measures and non-parametric tests were used for analysis comparing network activity. Paired t-tests were used for comparing within-subject behavioral response to optogenetic stimulation (two-tailed) or CNO application (one-tailed). We used a false discovery rate corrected alpha threshold for the three behavioral comparisons performed in each condition (aggressive behavior towards an intact male, non-aggressive interactions with an intact male, and non-aggressive interactions with a female). This correction was achieved using the Benjamini-Hochberg procedure. Because network activity is not normally distributed, we used a Friedman’s tests to compare repeated measures of network activity. Post-hoc testing was performed with two-tailed Wilcoxon signed-rank tests to compare within-subject activity, and two-tailed rank sum testing was used to perform between-subject network activity. This analysis approach was used to compare network activity responses to optogenetic stimulation, stimulus mouse exposure and interaction, and CNO injection. We used directional tests for our validation experiments because our initial dCSFA-NMF model mapped low *EN-AggINH* activity to aggression. Thus, we used one-tailed tests to determine whether manipulations known to increase aggression would also decrease network activity, and whether manipulations known to decrease aggression would also increase network activity. We used a Spearman rank to determine the relationship between network activity and aggressive behavior, and a Z test was used to determine if mice showed a greater proportion of cells that were coupled to network activity than would be expected by chance. We used a mixed effects model ANOVA to determine the impact of chronic mPFC → NAc LinCx-editing on behavior, and we used a false discovery rate-corrected alpha threshold to account for the three behavioral conditions. Data is presented as mean ± standard error of measurement, throughout the paper, unless otherwise specified.

## Data availability

Data will be made available for replication purposes and pre-agreed upon scientific extensions with a material transfer agreement. Please send all requests to Kafui Dzirasa or David Carlson. Responses will be provided within two weeks. Source data are provided with this manuscript.

## Code availability

This learning algorithm is publicly available code found at https://doi.org/10.7924/r4r578. Data will be made available for replication purposes and pre-agreed upon scientific extensions with a material transfer agreement.

## ACKNOWLEDGEMENTS

We would like to thank Stephen Mague, Karim Abdelaal, and Ashleigh Rawls comments on this work; Jean Mary Zarate for substantial guidance on revising this work; and Timothy Nyangacha for technical support.

## AUTHOR CONTRIBUTIONS

Conceptualization and Methodology – YSG, AT, NMG, SJR, DEC, and KD; Formal Analysis – YSG, AT, NMG, AFS, KKW-C, DEC, and KD; Investigation – YSG, AT, NMG, GET, AFS, KKW-C, SJR, DEC, and KD; Resources – YSG, AT, NMG, GET, AFS, KKW-C, SJR, DEC, KD; Writing – Original Draft, YSG, AT, NMG, SJR, DEC, and KD; Writing – Review & Editing, YSG, AT, NMG, KKW-C, SJR, DEC, and KD; Visualization – YSG, AT, DEC, and KD; Supervision – KKW-C, DEC, and KD; Project Administration and Funding Acquisition – SJR, DEC, and KD; See Supplementary materials for detailed author contributions.

