## Supplementary Materials for "Precision editing of an aggression-encoding network relay suppresses violent action in mice"

### SUPPLEMENTARY TABLE S1: DETAILED AUTHOR CONTRIBUTIONS

| Author | Contribution |
|---|---|
| **Yael S. Grossman** | Conceptualized and designed all experiments with SJR, DEC, and KD; Developed analytical strategy to discover network selectively encoding aggression with AT, DEC, and KD; determined brain region targets; designed electrodes for implantation; modified closed-loop stimulation code developed by NMG; assembled all electrodes and optrodes used for experiments; performed all viral and electrode implantation surgeries; bred ESR1 mice; castrated male C57 mice; collected all neurophysiological and behavioral data; performed all histological analysis; manually labelled all video data used to train initial aggression model and to perform the first network validation experiments in hold-out CD1 mice; performed automated video analysis of behavior for all optogenetic and DREADD manipulation experiments; performed all LFP data processing and LFP feature generation; determined network scores for all the hold-out testing animals/experiments; determined network activity stimulation threshold for closed-loop stimulation experiment; performed all statistical analysis of behavior and network activity for all validation experiments; performed all cell firing vs. electome network activity correlation analysis; designed/built (with KKW-C) and implanted electrodes (with KD) for dual micro-wire/silicon probe recordings; created all main figures with KD; Wrote original draft of the manuscript with AT, NMG, and DEC; reviewed and edited manuscript. |
| **Austin Talbot** | Converted published dCSFA-NMF model to tensor flow to 2.0 (Mague et al., 2022; Talbot et al., 2020); Trained all statistical models utilized in the study; determined hyperparameters for discovering the final aggression network (including the number of total networks to including in training); developed and validated statistical approaches to discover the aggression network, including using individual behavioral windows pooled across mice to train models; created supplementary figure outlining hyperparameter selection for network training; wrote the original draft with YSG, NMG, and DEC; reviewed and edited the manuscript. |
| **Neil M. Gallagher** | Developed feature generation pipeline; developed methods for closed-loop stimulation including optimization of real-time feature calculation; developed method for nonsynchronous stimulation; conceptualized strategy for established mouse-specific network stimulation threshold for closed-loop manipulation; conceptualized behavioral protocol for evaluating impact of closed loop stimulation; Assisted with performing a subset of recording and stimulation experiments to |

| | |
|---|---|
| | validate closed-loop methodology; analyzed final closed-loop stimulation results; wrote the original draft with YSG, AT, and NMG; methodological description of closed-loop stimulation experiments; reviewed and edited the manuscript. |
| **Kathryn Katsue Walder-Christensen** | Trained DeepLabCut (DLC) and Simple Behavioral Analysis (SimBA) models to automatically classify social behaviors during aggressive and control encounters; Reviewed and edited manuscript; Assisted KD with supervision of all behavioral experiments. Designed electrodes and implantation procedure for concurrent microwire/silicon probe recordings with YSG; Performed high density cellular recordings with YSG. Performed histology for high-density cellular recording experiment and LinCx experiment with YSG; reviewed the manuscript. |
| **Gwenaëlle E. Thomas** | Provided critical optogenetics training to YSG; Assisted with optogenetics behavioral experiments; reviewed the manuscript. |
| **Alexandra Fink Skular** | Sorted cellular action potentials for subsequent performed cell firing vs. electome network activity correlation analysis; verified baseline CD1 aggression levels for a subset of experiments; reviewed the manuscript. |
| **Scott J. Russo** | Conceptualized protocol for social behavioral testing in CD1 mice with YSG and KD; conceptualized approach for inducing aggression toward female mice using ESR1-Cre mice; conceptualized optogenetic stimulation approach utilized for ESR1-Cre mice; provided ESR1-Cre mice; revised the original draft of the manuscript; reviewed and edited paper; secured funding for the study. |
| **David E. Carlson** | Conceptualized feature generation with NMG; conceptualized and supervised methodology and code implementations for closed-loop stimulation with KD and NMG; conceptualized visualizations; wrote original draft with YSG, AT, and NMG; reviewed and edited paper; supervised all machine learning model development and analyses; provided and maintained computational resources for machine learning implementation; secured funding for the study. |
| **Kafui Dzirasa** | Conceptualized general approach to mapping networks that discriminate between distinct social behaviors with DEC; conceptualized approach to closed-loop optogenetic stimulation based on electome network activity with NMG and DEC; conceptualized approach to analyze encoding of social behaviors based on activity within single brain regions; implanted electrodes for dual micro-wire/silicon probe recordings with YSG; created main figures with YSG and DEC; substantially revised the original draft of the manuscript; reviewed and edited paper; supervised all behavioral experiments with assistance of KKW-C, supervised analysis of behavioral |

| | |
|---|---|
| | and neurophysiological results with DEC; secured funding for the study. |
| ****<u>Attribution Process</u>** | Each team member outlined their individual contributions across a standard set of domains (conceptualization and methodology, formal analysis, investigation, resources, writing -original draft, writing -review & editing, visualization, Supervision, and Project Administration and Funding Acquisition) and subsequently had the opportunity to edit a summarized attribution description to their satisfaction. Contribution summaries were then shared across all team members. Each team member had the opportunity to raise concerns with regard to any other team member's outlined contributions, and issues that remained unaddressed after additional revisions were subjected to a mediation process led by the lead principal investigator. The assigned authorships and these detailed author contribution descriptions reflect the outcome of this process. |

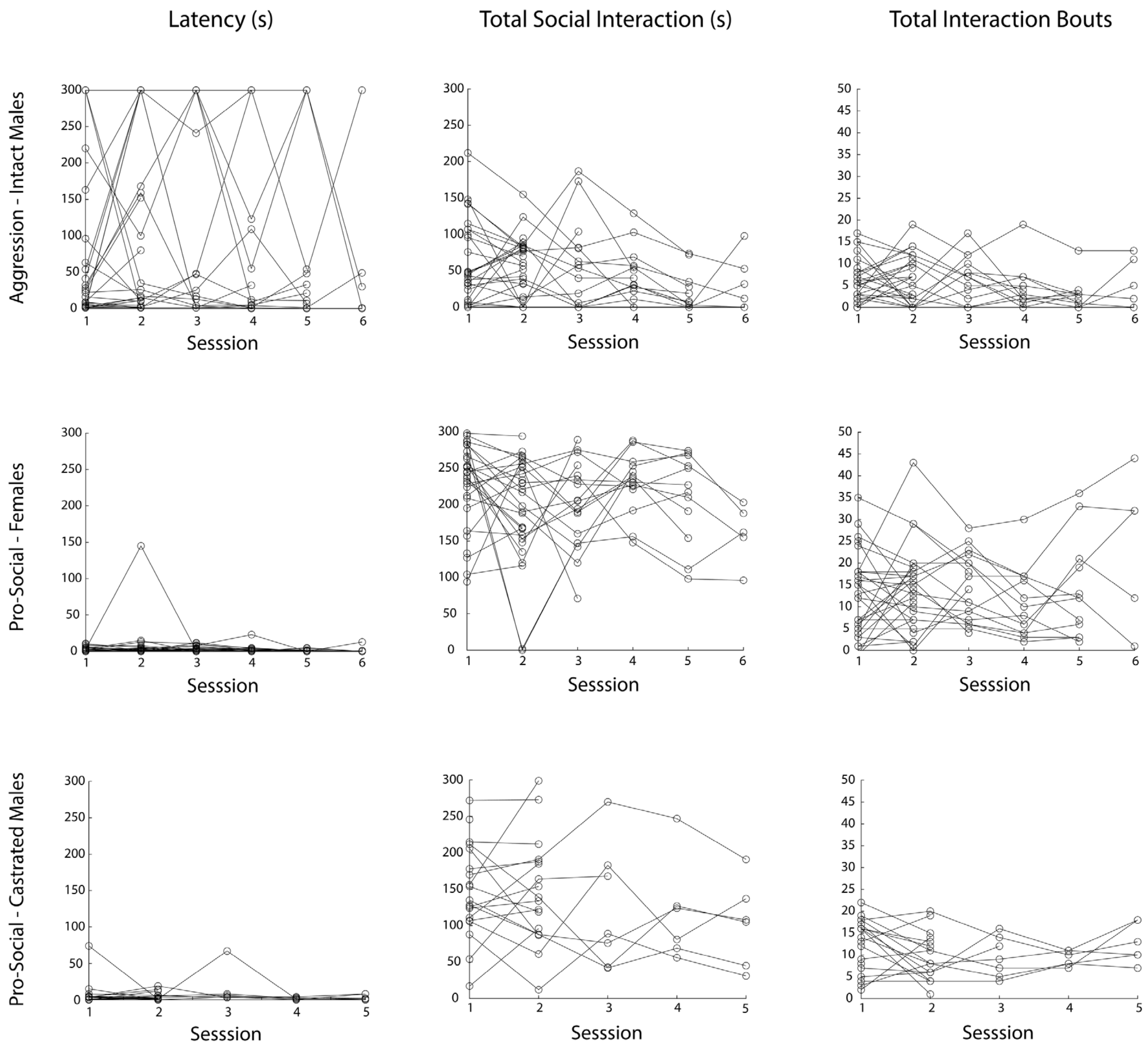


**Supplementary Figure S1: Aggressive and non-aggressive behavior across recording sessions.** The latency of CD1 mice to engage in social interaction (left column), the total social interaction time (middle column), and the number of interaction bouts (right column) are shown for aggressive sessions with intact males (top row), and non-aggressive sessions with females (middle row) and castrated males (bottom row). N=31 CD1 mice.

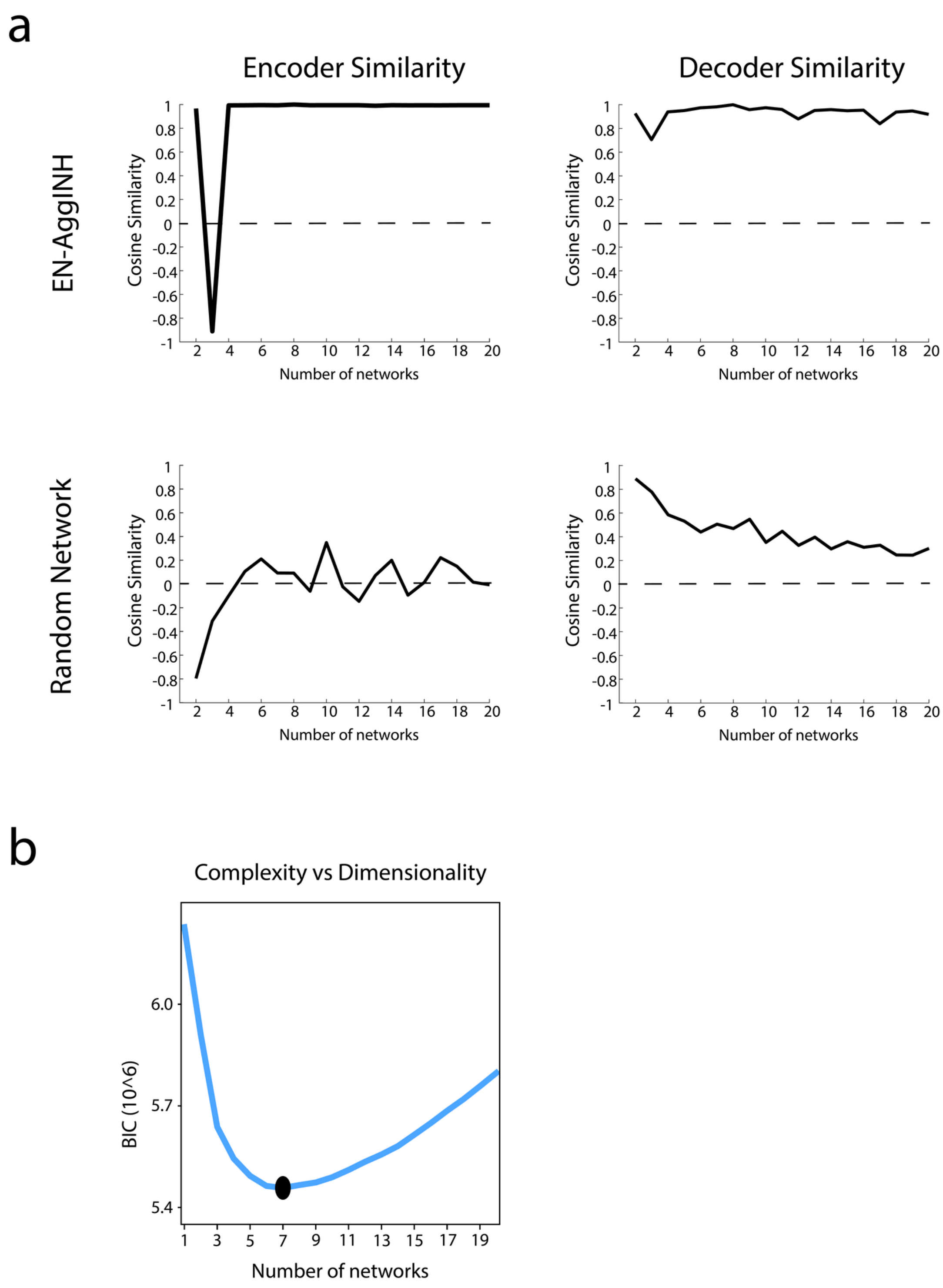


**Supplementary Figure S2: Validating Model Dimensionality. a)** Top: The cosine similarity between *EN-AggINH* as compared to the supervised electome in models learned with different numbers of networks. Cosine similarity is calculated between two vectors $a$, $b$ as $a^T b / \|a\|_2 \|b\|_2$, where a value of 1 denotes the vectors are in the exact same direction from the origin and a value of -1 denotes that the vectors are in the exact opposite directions. The supervised encoder is highly stable across models with varying dimensionality (left). The only exception is the model employing three networks, which learned a positively associated aggression network rather than the negatively associated network recovered in all other models; however, this difference reflects only an inversion in directionality and is otherwise similar, as sign is adjusted for in the downstream classifier. Across models containing 5–20 total networks, cosine similarity values for both the supervised encoder and decoder exceed 0.95 relative to the final model, demonstrating that the aggression-related latent factor and its mapping to behavior are robust to the choice of dimensionality within this range. Bottom: To contextualize the magnitude of the cosine similarity values observed for the supervised network, we computed cosine similarity between randomly selected unsupervised networks across models. Generative encoders for unsupervised networks show little similarity when compared across models (left), and their corresponding network decoders likewise do not exhibit high cosine similarity (right). This comparison demonstrates that high cosine similarity values obtained for the supervised network are not trivially obtained across models and supports the conclusion that the supervised aggression network is uniquely stable across dimensionalities. **b)** We utilized the Bayesian Information Criterion (BIC) to select the number of unsupervised networks to use in the final network model. The BIC was evaluated for all dimensionalities from 1-20 factors, and the lowest value of 7 was selected as the best model. With the supervised network, this means we use eight networks in total.

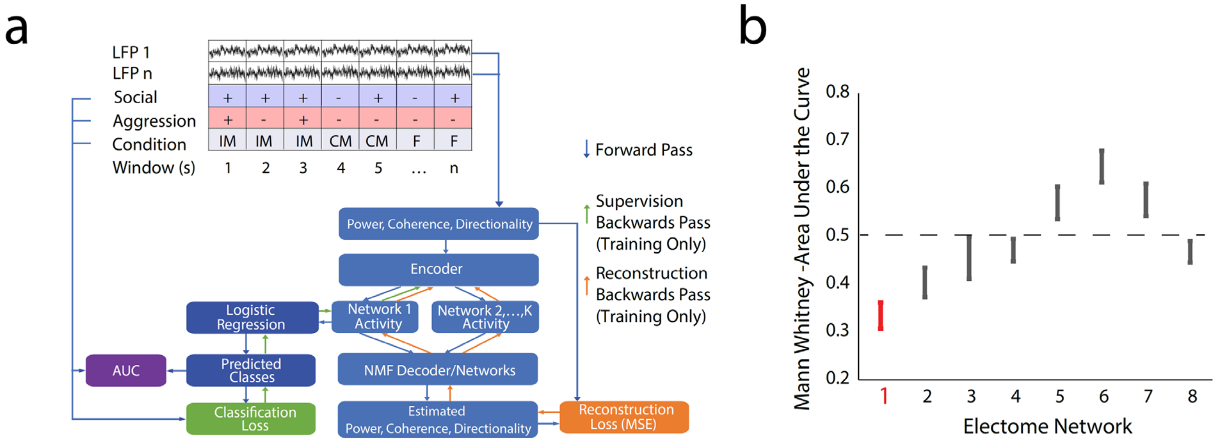


**Supplementary Figure S3: dCSFA-NMF model and network training outcomes. a)** Schematic of machine-learning model used to discover network encoding aggressive behavior. (Top) The LFP data recorded concurrently from microwires implanted across eleven brain regions were segmented into 1-second windows and aligned with the behavioral classifiers and test conditions for those windows. The behavioral classifiers indicated whether animals were engaged in social behavior (+/-; light blue highlight), and if so, whether that social behavior was aggressive or not (+/-; pink highlight). The test conditions included intact male intruders (IM), castrated male intruders (CM), and female intruders (F). (Bottom) The LFP data for each window was used to calculate power, coherence, and directionality features across frequencies of 1-56Hz. These data were the inputs to the model. When inferring the network activities and predictions, the data flows along the forward pass arrows, where the features are extracted from the LFP data and then passed to the encoder to estimate all network activities. From there, the first network's activity is used in the logistic regression to predict whether the animal is in an aggressive state or not. All network activities are then multiplied by their corresponding learned network weight vectors (i.e., the NMF decoder) and added together to reconstruct the estimated neural feature matrix comprising power, coherence, and directionality. During training, the gradients flow along the backwards pass arrows, where all networks and activities are influenced by the ability to reconstruct the neural data, but only the Network 1 activity is directly impacted by the classification loss (light green). The collected gradients on the neural activities then continue backwards to learn the encoder parameters. This creates a full system that is learned by stochastic optimization and can efficiently estimate network activities through the encoder after training. **b)** Mann-Whitney Area Under the Curves (MW-AUCs) are shown for data from hold-out mice (N=9 mice). A MW-AUC of 1 indicates that high network activity predicts that animals are engaged in aggressive behavior towards intact males. Conversely, a MW-AUC of 0 indicates that low network activity predicts that animals are engaged in aggressive behavior, and a MW-AUC of 0.5 indicates that network activity does not predict aggressive vs. non-aggressive social behavior. Low Network 1 activity (the supervised network, highlighted in red) predicted aggressive behavior, while high Network 6 activity predicted aggressive behavior. The edges of the lines for each network extend to the mean±s.e.m. Source data are provided as a Source Data file.

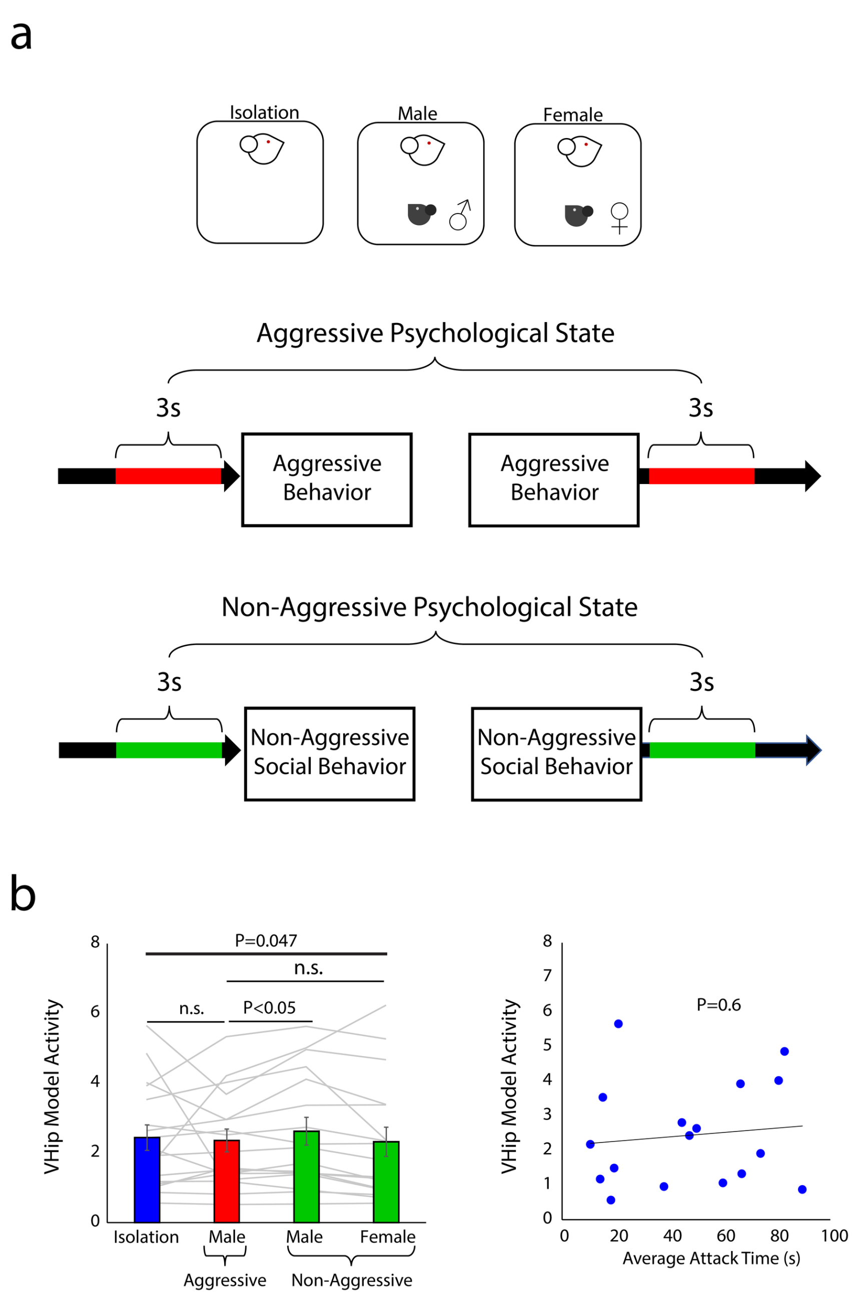


**Supplementary Figure S4: Ventral hippocampus (VHip) activity does not encode a generalized aggressive psychological state. a)** Neural activity was sampled while 17 mice were socially isolated (blue) and during intervals preceding and following their aggressive behavior towards intact male mice (red) and non-aggressive social behavior towards intact male and female mice (green). These data windows were not used to train the network model since they did not contain the behaviors of interest. **b)** Activity from ventral hippocampus was statistically indistinguishable between periods surrounding aggressive behavior towards intact males and non-aggressive social interactions with females and activity during social isolation ($F_{3,67}$=7.9, P=0.047 using Friedman's test; P=0.36 and 0.72, respectively, using post-hoc two-tailed Wilcoxon signed-rank test, significance determined by FDR correction, N=17 mice from the training set; Fig. 2b, left). No relationship was found between ventral hippocampal activity during isolation and the innate aggressiveness of mice (P=0.60 using two-tailed Spearman's rank correlation, Fig. 2b, right). Bar graphs show mean, and error bars show mean±s.e.m. Source data are provided as a Source Data file.

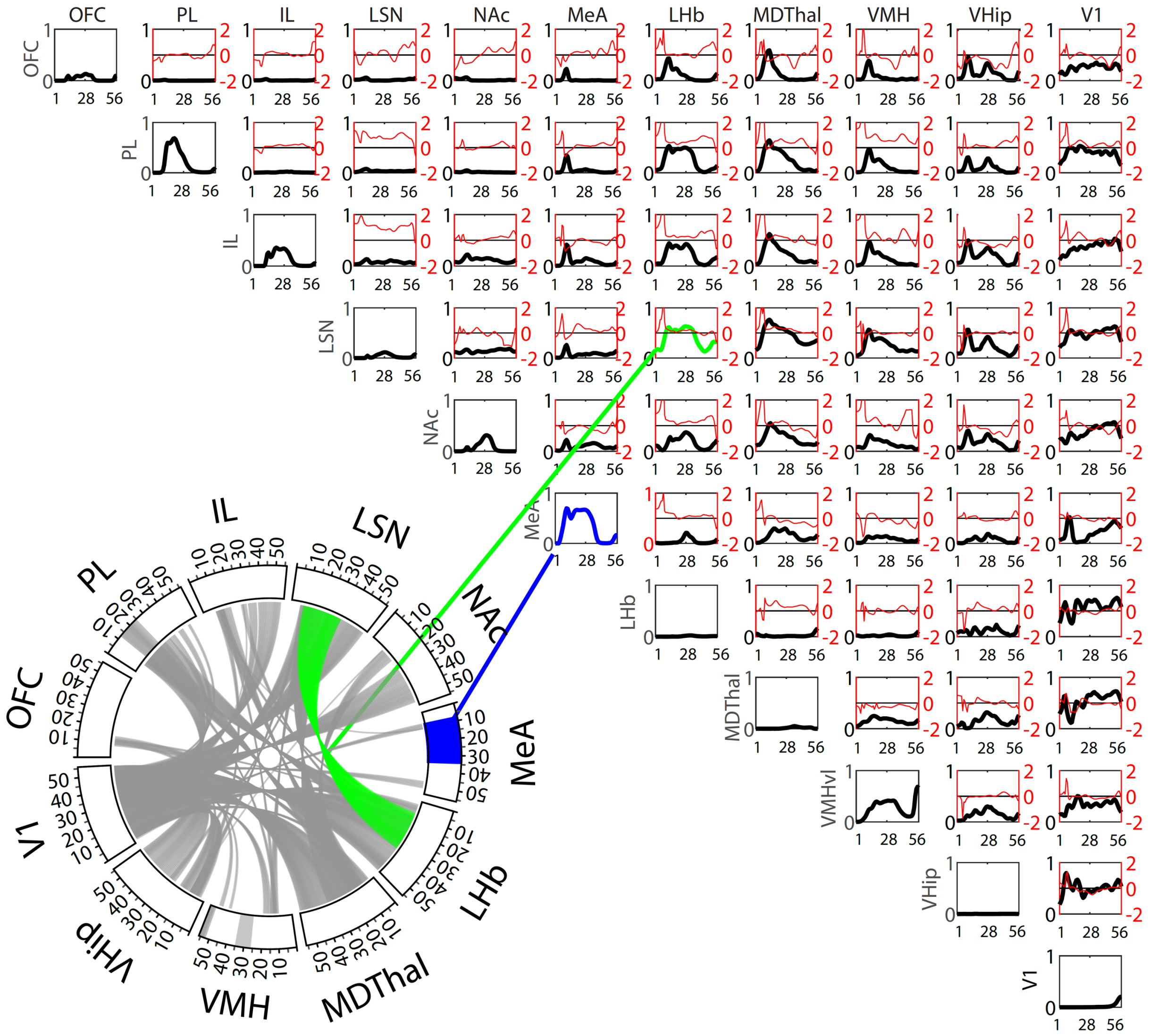


**Supplementary Figure S5: Power and coherence measures that define *Electome Network 1* (*EN-AggINH*).** Brain areas are shown to the top and the left identifying power and coherence density functions for *EN-AggINH*. Amplitude values (shown in black) reflect the relative LFP spectral energy (RLE) observed at each frequency, where the *Electome Network* is normalized to the total energy observed across the 8 networks (i.e., 1 supervised and 7 unsupervised networks). The offset between the two non-normalized directionality features for each brain area pair (A → B and B → A) are also shown in red (axis scale to the right). Positive spectral offsets correspond to frequencies at which the area listed along the top leads the area listed on the left. Negative spectral offsets correspond to the frequencies at which the area listed on the left leads the area listed on the top (Mague *et al.*, 2022). The circular plot depicts the frequencies for power (outer rim) and coherence (curved lines connecting two regions) above an amplitude threshold of 0.46, corresponding to the top 10% of predictor weights. As a representative example, the power measures for medial amygdala are highlighted in blue in both the circular and correlation plots; coherence between lateral septum and lateral habenula is highlighted in green.

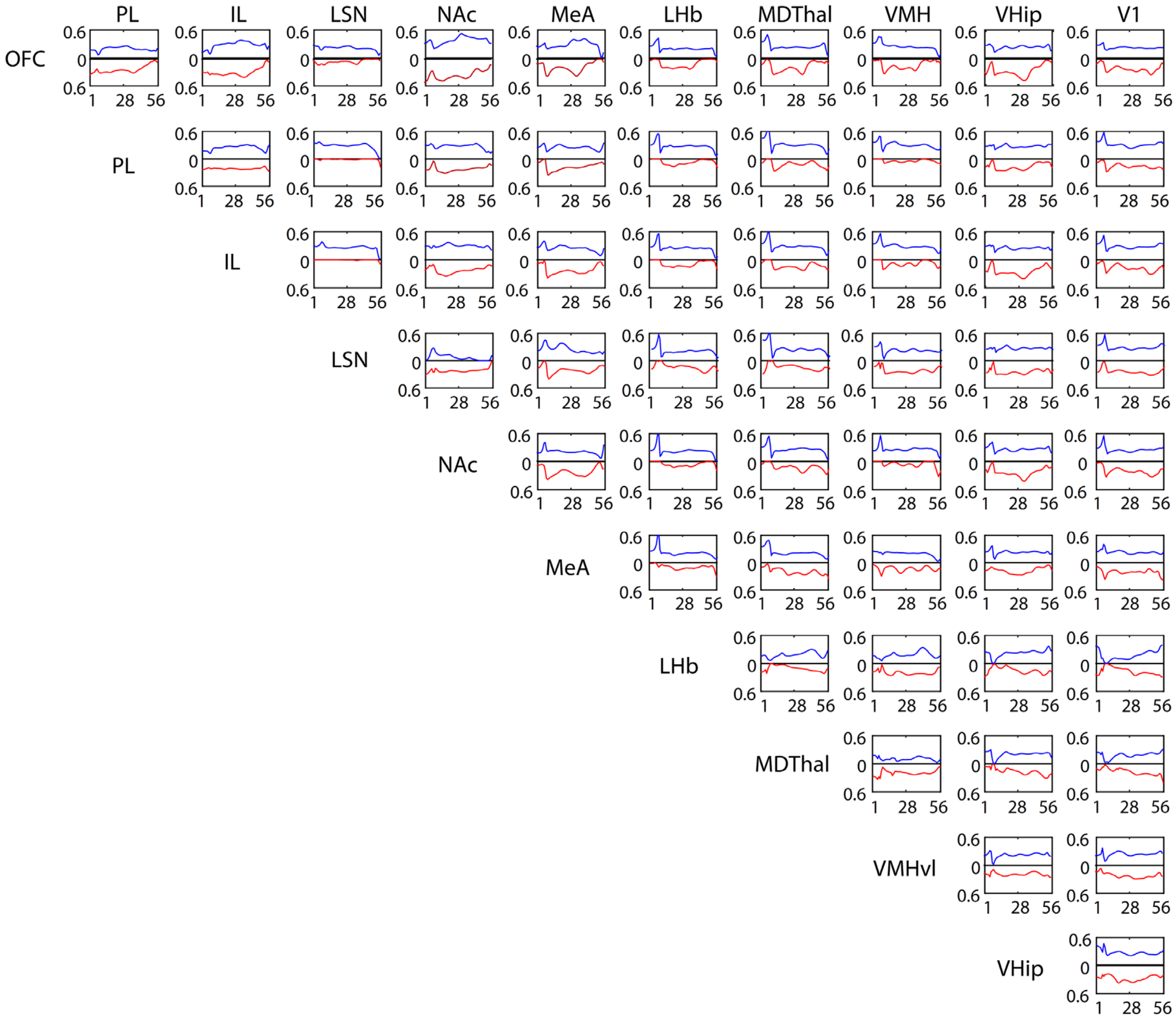


**Supplementary Figure S6: Directionality features that define *Electome Network 1* (*EN-AggINH*).** Data depicts the relative LFP spectral energy (RLE) observed at each frequency, where the network is normalized to the total energy observed across the 8 networks (i.e., 1 supervised and 7 unsupervised networks). Red lines correspond to directionality from the area on the left to the area on the top (0 to 0.6 RLE; note that these plots are inverted), while the blue lines correspond to the directionality from the area on the top to the area on the left (0 to 0.6 RLE).

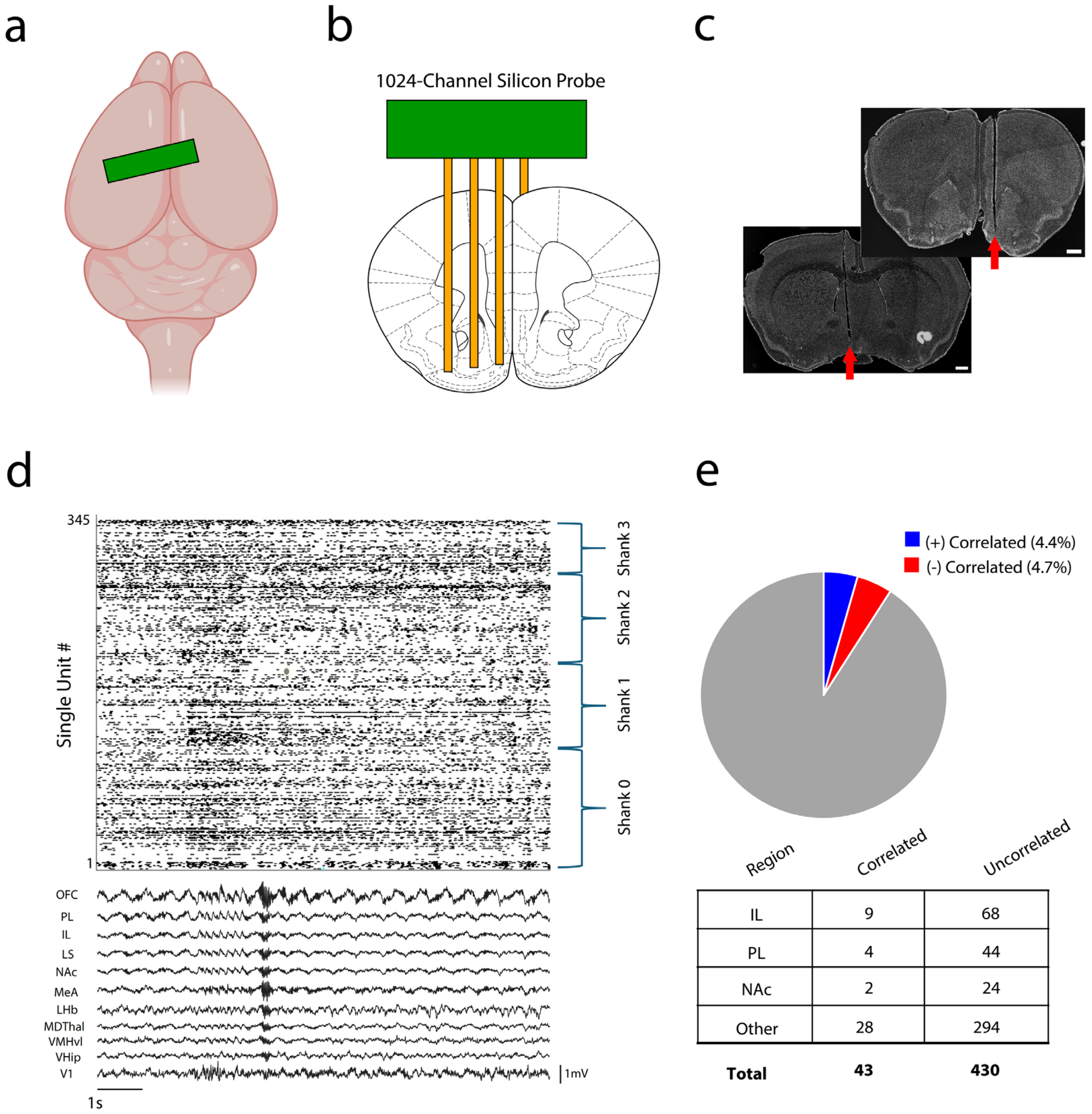


**Supplementary Figure S7: High-density recordings to quantify coupling between cellular and network activity. a-b)** Target sites for 1024-channel silicon probe implantation. The green rectangle in (a) corresponds to the silicon probe placement. **c)** Representative histological images showing tracks of silicon probes (red arrows; images from one of the two implanted mice, scale bars correspond to 500µm). **d)** Raster plot showing 345 single units recorded concurrently with LFPs from the 11 brain regions that compose *EN-AggINH*. **e)** Portion of single units recorded from two CD1 mice which showed activity that was correlated (blue) or anticorrelated (red) with *EN-AggINH* activity (top). The number of a region's single units whose activity was correlated or uncorrelated with *EN-AggINH* activity is shown on the bottom. Note that counts are shown for mPFC (PL and IL) and NAc, the regions that compose the circuit targeted for LinCx editing. Source data are provided as a Source Data file.

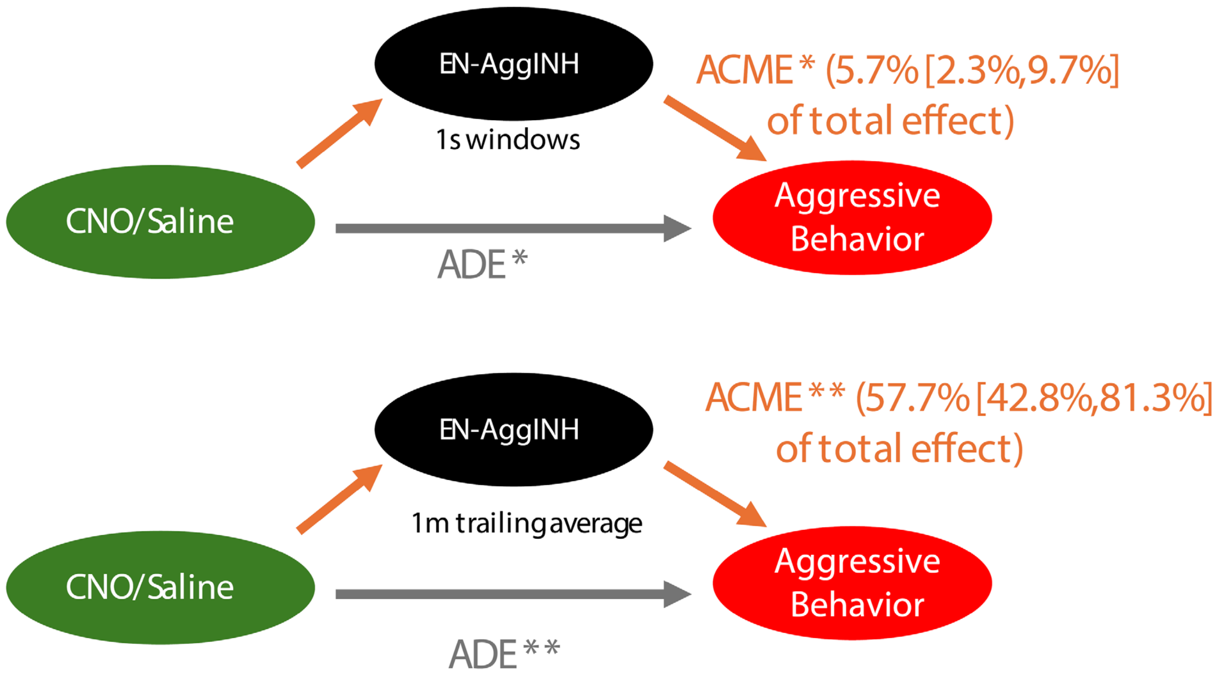


**Supplementary Figure S8:** ***EN-Aggression Inhibition (EN-AggINH)* mediates aggression in response to causal activation of ESR1+ cells in VMHvl.** We considered the CNO condition as the treatment case and saline as the control case and then used the same mediation approach as in the closed-loop stimulation. Network expression was considered as a potential mediator, and the outcome condition was aggressive interaction versus both non-aggressive interaction and non-interaction. Models were corrected for mouse identity as a mixed model. Each second of data was considered as a separate sample. In this case, both the average causal mediated effect (ACME) and the average direct effect (ADE) were significant (*P<0.01). However, the relative fraction of behavioral change caused by the treatment explained by the ACME was relatively small (5.7%). We attribute this due to the challenges of the long-term impact of the treatment intervention, which changes the overall emotional state of the animal rather than just on a second-to-second basis. As such, we evaluated what would happen if we used the trailing average of *EN-AggINH* over the last minute rather than the instantaneous value to account for the fact that the treatment is changing behavior over a longer time frame. By using this average value of *EN-AggINH*, we found that 57.0% (**P<0.0001) of the treatment effect of CNO was mediated by the network expression. This result suggests the *EN-AggINH* does in fact significantly mediate behavioral response to treatment in multiple interventional experiments, further strengthening our prior conclusions.

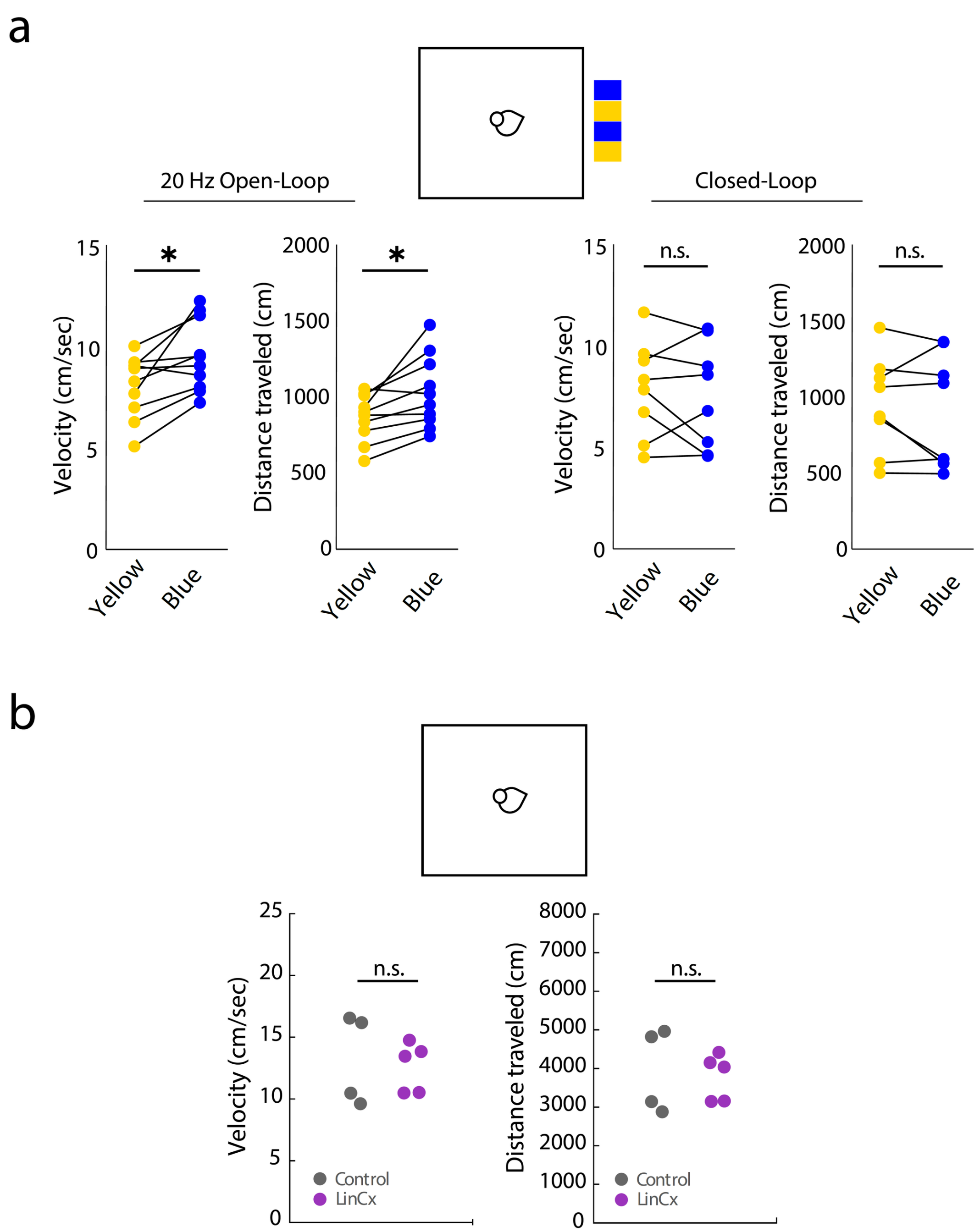


**Supplementary Figure S9. Gross locomotor impact of PFC circuit modulation. a):** Off-target effects of open-loop PFC stimulation. Mice were tested across four segments of alternating blue or yellow light stimulation; order of stimulation segments is shown at the top. Each segment was 2 minutes long. Open-loop stimulation of prefrontal cortex with blue light at 20Hz induced behavioral hyperactivity in the CD1 mice (N=10; $T_9$= 3.221 and *P = 0.011 for Velocity; $T_9$= 3.213 and *P=0.011 for distance traveled, using two-tailed paired t-test for blue vs. yellow light, left). On the other hand, closed-loop stimulation of prefrontal cortex with blue light had no impact on gross locomotor behavior (N=8; $T_7$= 0.58 and P=0.58 for velocity; $T_7$= 0.86 and P= 0.42 for distance traveled, using two-tailed paired t-test for blue vs. yellow light; right). **b)** LinCx-editing the mPFC → NAc circuit had no impact on gross locomotor behavior (N=4 and 5 for control and LinCx-edited mice, respectively; $T_7$= 0.31 and P = 0.77 for Velocity, left; $T_7$= 0.30 and P=0.77 for distance traveled, using two-tailed paired t-test, right). Mice were tested for 5 minutes. Source data are provided as a Source Data file.

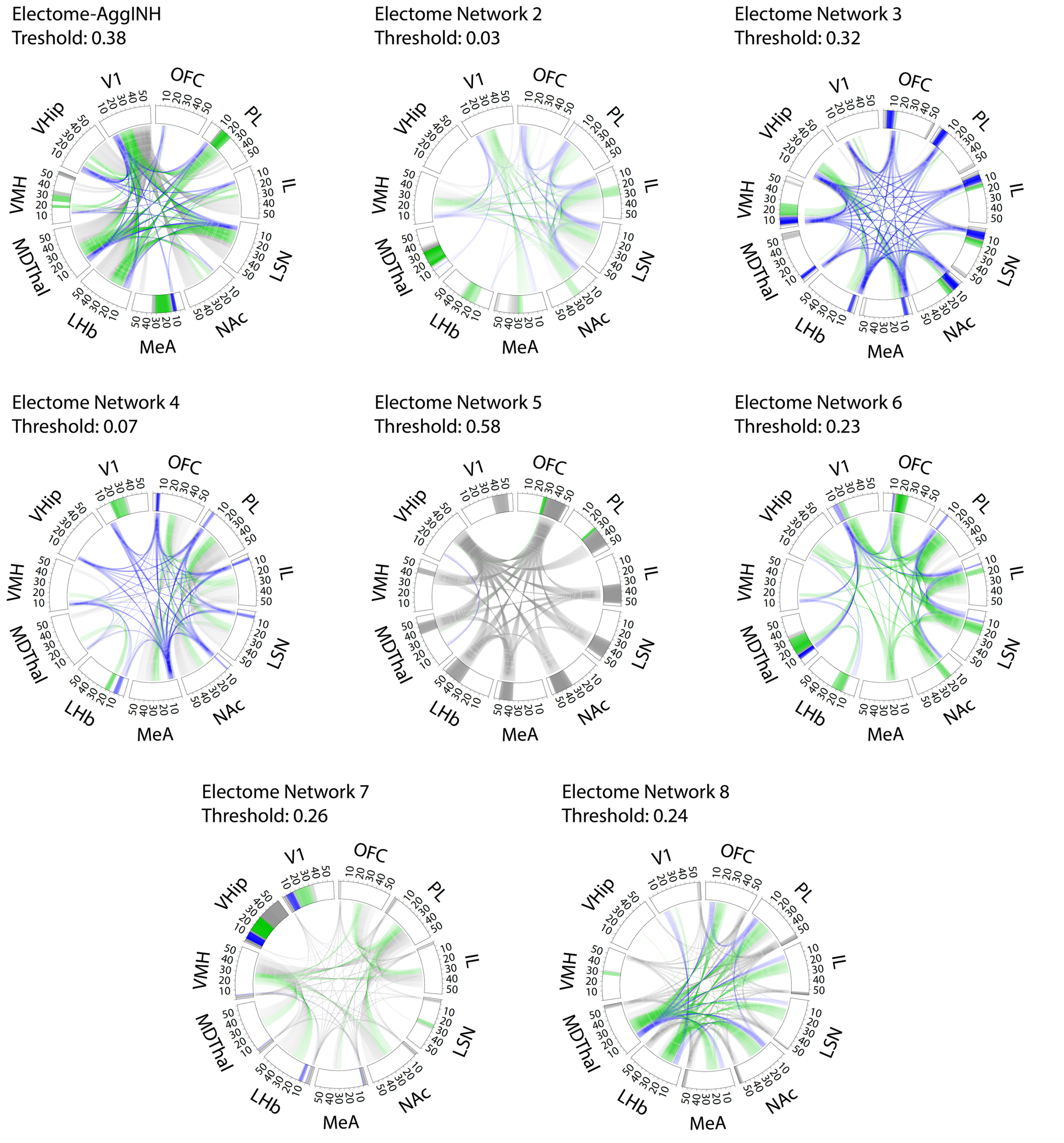


**Supplementary Figure S10. Components of the eight learned *Electome Networks* (*EN-AggINH* and the 7 unsupervised networks)*.** *Prominent oscillatory frequency bands highlighted for each brain region around the rim of the circle plot. Prominent coherence measures are depicted by lines connecting brain regions through the center of the circle. Each plot is shown at relative spectral density corresponding to the top 85% of features for that network. Theta (4-11Hz) and beta (14-30Hz) frequency components are highlighted in blue and green, respectively. Networks 4-7 have strong contributions of primary visual cortex (V1) power and Network 7 has strong contributions of ventral hippocampal (VHip) power.

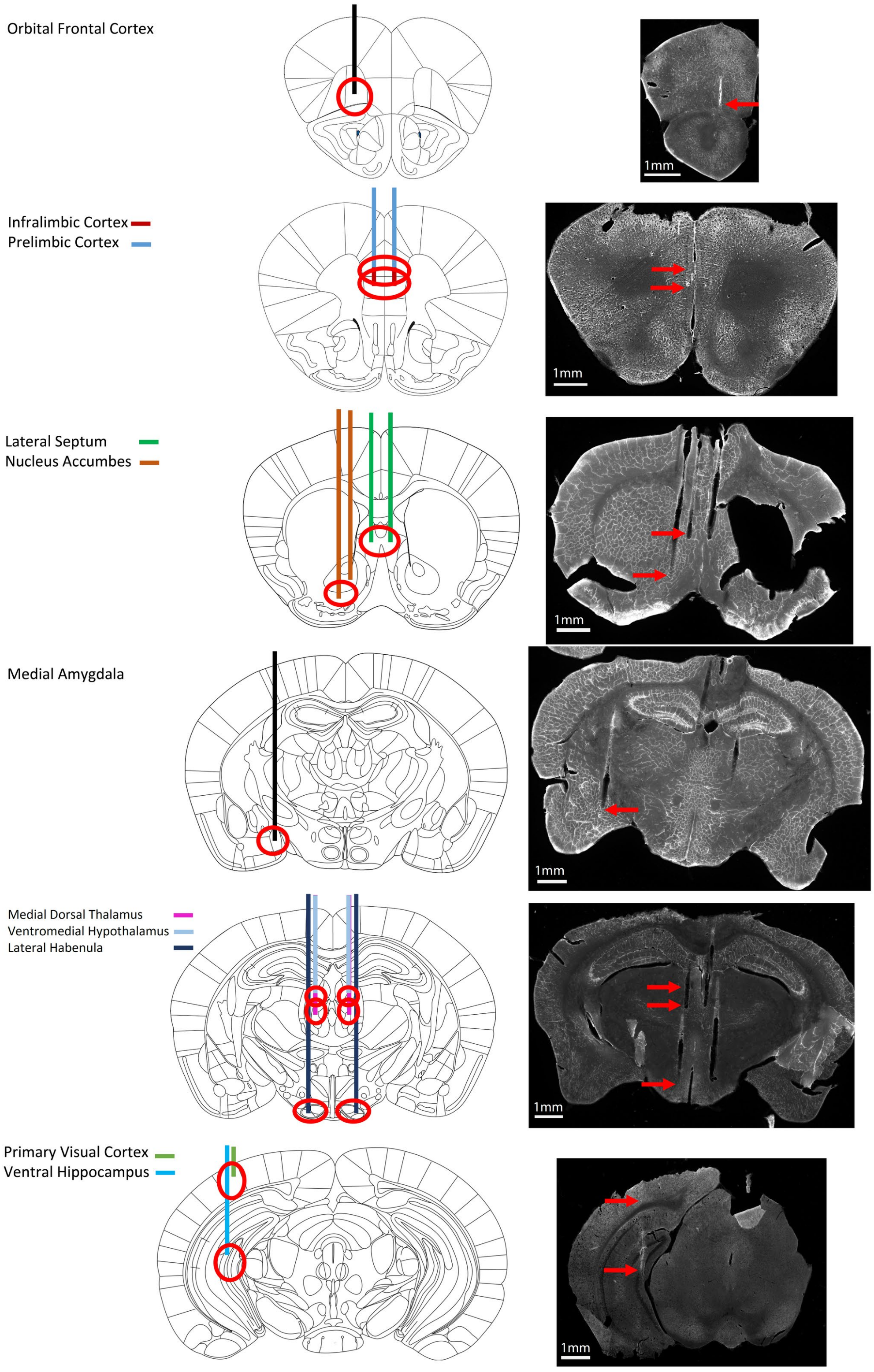


**Supplementary Figure S11: Electrode targeting strategy and histological confirmation.** Electrode bundles were centered within the red circles shown for each target brain area (left). For brain slices depicting more than one target region, the color of the electrode corresponds to the region listed to the left. Representative histological images are shown with red arrows highlighting electrode tracks (right).

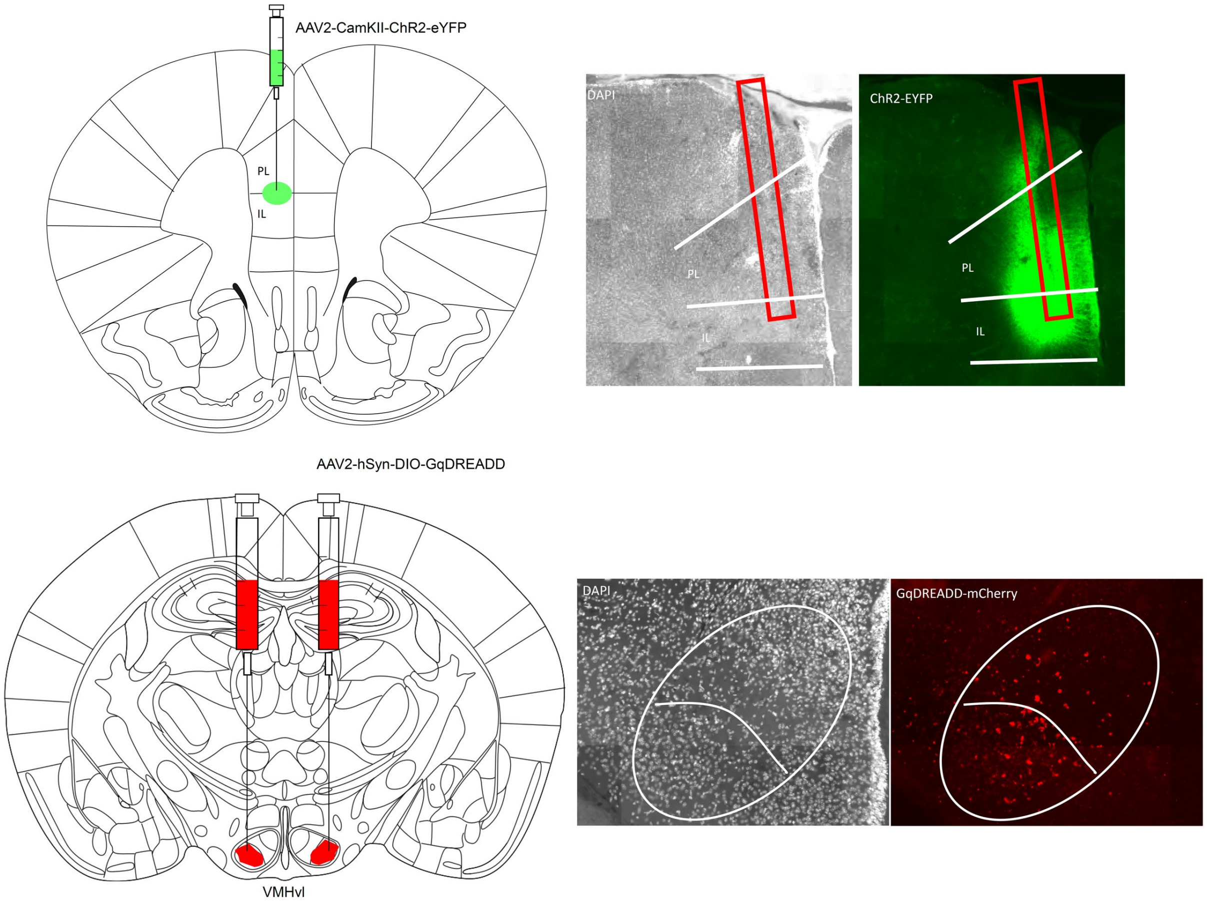


**Supplementary Figure S12: Representative histological images showing targeting and expression of ChR2 and Gq-DREADD for single-region manipulations.** Top: The white lines in the right image highlight the anatomical boundaries of PL and IL, which are both targeted as depicted in green in the left image. Red rectangles correspond to the insertion tracks for the optic fibers. Bottom: The white ellipse highlights VMH, and the ventrolateral nucleus is demarcated below the curved line (targeted and labeled in red in the left image).

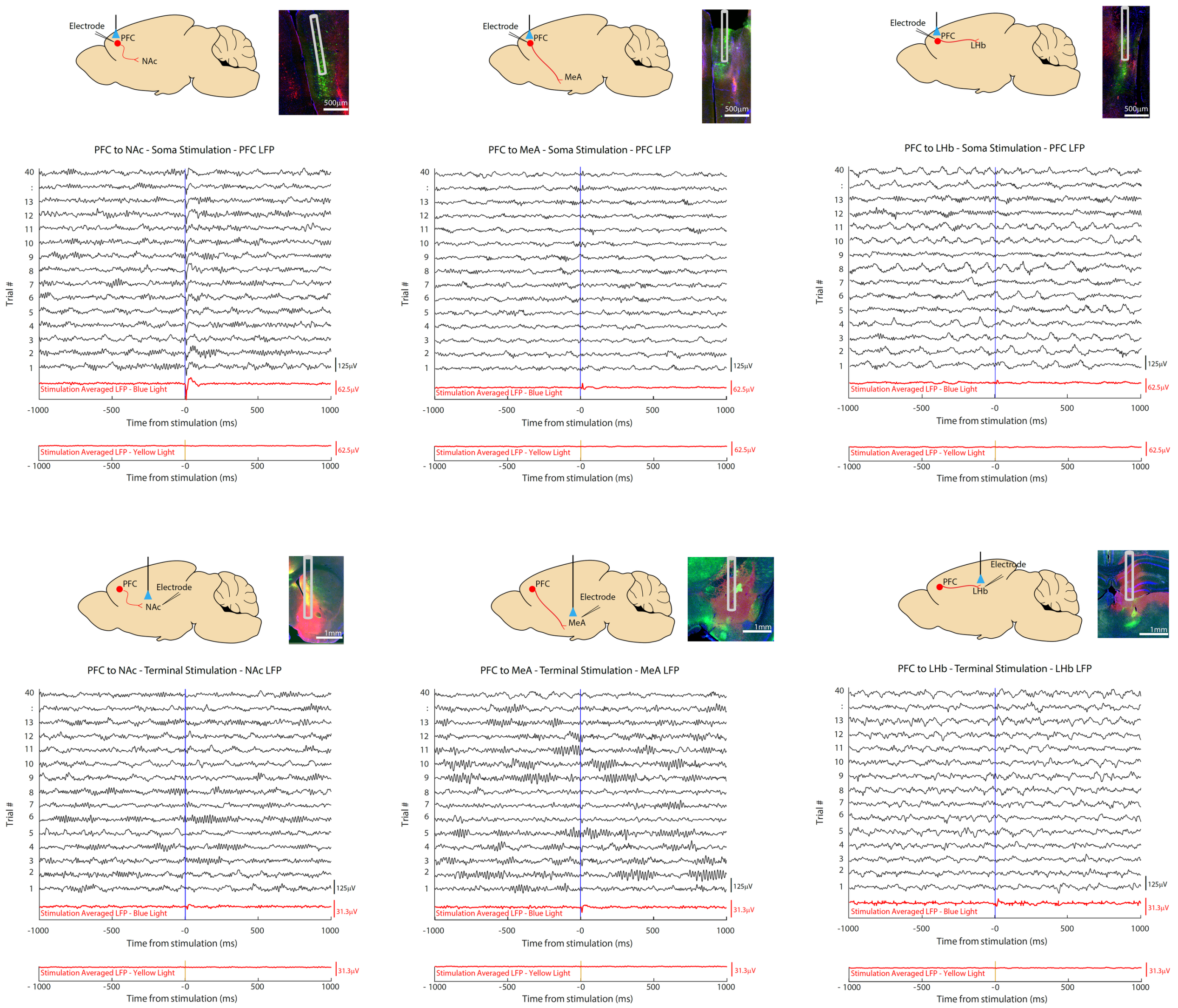


**Supplementary Figure S13: Representative histological images showing targeting and expression of ChR2 and retrograde Cre AAVs.** The green fluorescence is indicative of the ChR2-EYFP construct, while the red fluorescence is indicative of the Cre-mCherry construct. Data are shown for each of three circuits. Each plot shows representative LFP oscillations in response to an optogenetic light pulse (blue vertical line; light stimulation was delivered at 5mW, 10ms pulse width). LFP activity averaged across 40 blue light pulses is shown below in red. For the top row, light was delivered to the prefrontal cortex (PFC), and LFP was recorded from the PFC. For the bottom row, light was delivered to the PFC terminals in one of the target regions (Nucleus Accumbens – NAc/left, Medial Amygdala – MeA/middle, Lateral Habenula – LHb/right), and neural activity was recorded from the same target region. Blue light stimulation induced neural activation at each targeted site ($P<0.05$ determined using bootstrapping at an $\alpha=0.05$, Bonferroni-corrected for 60 comparisons). Potentials evoked by yellow light are shown on the bottom of each plot. No significant evoked potentials were observed for any of the circuits in response to yellow light stimulation. Scale bar = 500μm for all histological images in the top row, 1mm for all histological images in the bottom row.

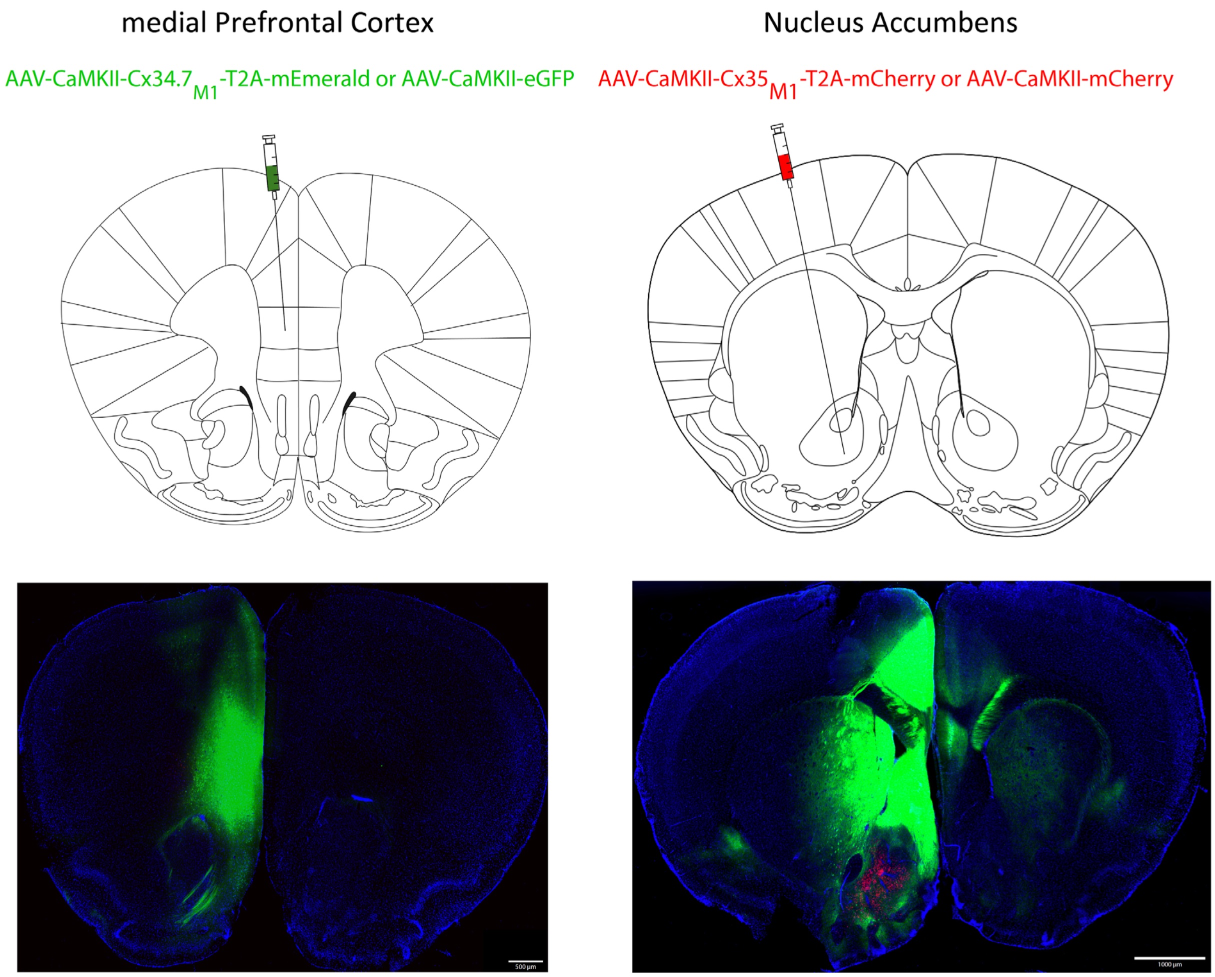


**Supplementary Figure S14. Representative histological images showing targeting and expression of LinCx and control viruses.** Tissue was collected 8 weeks post-surgery. Images on the left depict expression of Cx34.7$_{M1}$-T2A-mEmerald at the medial prefrontal cortex target site (green). Images on the right depict NAc- targeted afferents from the medial prefrontal cortex (mPFC) that are observable at this time. Expression of Cx35M1-T2A-mCherry is also shown in the NAc (red). Note the overlap between the green afferents and red-tagged soma.
